# RNA nucleotide repeats induce mitochondrial dysfunction and the ribosome-associated quality control

**DOI:** 10.1101/2021.09.24.461665

**Authors:** Joana Teixeira, Anu-Mari Harju, Alaa Othman, Ove Eriksson, Ana Brandão, Brendan J. Battersby, Susana M. D. A. Garcia

## Abstract

Nucleotide repeat sequences are prevalent in the genome and expansion of these sequences is associated with more than 40 neuromuscular disorders. To understand the pathogenic mechanisms underlying RNA-repeat toxicity, we performed a genetic screen in a *Caenorhabditis elegans* model expressing an expanded CUG repeat specifically in the muscle. Here, we show that expression of this RNA repeat impairs motility by mitochondrial dysfunction, disrupting mitochondrial morphology and respiration. The phenotype is dependent on the RNA-binding factor MBL-1 and requires factors from the ribosome-associated protein quality control complex. Furthermore, Coenzyme Q supplementation rescued the motility impairment and all of the mitochondrial phenotypes. Together, our data reveal the importance of mitochondrial dysfunction in the molecular pathogenesis of RNA repeat expansion disorders.

## Introduction

Nucleotide repeat sequences are prevalent in the coding and non-coding regions of the genome(Lander et al., 2001). These sequences can form diverse secondary structures and thus are prone to breakage, replication stalling and genome rearrangements that during DNA repair lead to repeat contractions or expansions(Brown and Freudenreich, 2021). The instability in repetitive sequences is associated with expansions in more than 40 neuromuscular disorders(Paulson, 2018). Expanded repeats often exhibit a genotype-phenotype correlation that affects disease severity and age of onset(Andrew et al., 1993; Tokgozoglu et al., 1995). At the cellular level, the pathogenic mechanisms of RNA repeat toxicity are complex: transcript loss-of-function; inappropriate interaction with RNA binding proteins(Miller et al., 2000); and non-AUG-dependent (RAN) translation of the repeats into toxic proteins(Swinnen et al., 2020). RNA repeats act as gain-of-function pathogenic agents by sequestering RNA binding proteins(Miller et al., 2000). Despite this, the underlying spectrum of mechanisms by which RNA repeats induce cellular dysfunction and contribute to the molecular pathogenesis remains poorly understood.

The most prevalent form of adult-onset muscular dystrophy(Harper, 2001), myotonic dystrophy type 1 (DM1), results from a CTG expansion in the 3’UTR of the dystrophia myotonica-protein kinase (DMPK) gene(Brook et al., 1992; Mahadevan et al., 1992). DM1 is a dominantly inherited systemic disorder characterized by progressive muscular weakness, atrophy and myotonia(Chau and Kalsotra, 2015; Harper, 2001). In patients, the CTG expansion ranges from 50 to 4,000 repeats, whereas healthy individuals have 5 to 35 repeats(Turner and Hilton-Jones, 2010). DM1 transcripts constitute the prototype gain-of-function RNA repeat toxicity, as any RNA bearing an expanded CUG repeat in the 3’ UTR is sufficient to induce DM1-like features(Mankodi et al., 2000). These transcripts can accumulate in the nucleus as discrete RNA foci^3^ and are predicted to affect the function of RNA-binding proteins. The best-characterized modifier of DM1 pathogenesis is the developmentally regulated RNA-binding factor family Muscleblind-like (MBNL)(Miller et al., 2000). The conserved MBNL family regulates many aspects of RNA metabolism, including alternative splicing(Ho et al., 2004; Wang et al., 2012), subcellular localization(Adereth et al., 2005; Wang et al., 2012), mRNA stability(Masuda et al., 2012; Wang et al., 2012) and alternative polyadenylation(Batra et al., 2015). Expanded RNA repeats sequester MBNL and affect these critical functions(Batra et al., 2014; Goodwin et al., 2015; Masuda et al., 2012; Wagner et al., 2016). In *Caenorhabditis elegans*, MBL-1 is the sole orthologue of this RNA binding protein, which also binds CUG repeats(Garcia et al., 2014; Sasagawa et al., 2009) and co-localizes with toxic RNAs(Garcia et al., 2014). As MBNL dysfunction has broad effects on the DM1 transcriptome, how these changes are linked to the pathogenesis is less understood.

To identify molecular factors that modulate RNA toxicity, we took advantage of an established *C. elegans* model with an expanded CUG repeat expressed only in the muscle to perform an RNA interference (RNAi) genetic screen. This model exhibits impaired motility and the accumulation of RNA ribonuclear foci associated with MBL-1, phenotypes observed in DM patients(Garcia et al., 2014). Here, we uncover a mechanism by which RNA repeat toxicity drives mitochondrial dysfunction involving MBL-1 and the ribosome-associated quality control complex (RQC). Metabolic supplementation with coenzyme Q (CoQ) rescued all of the mitochondrial phenotypes and the movement defect. Together, these findings point to the importance of mitochondrial dysfunction and cytosolic protein homeostasis in the molecular pathogenesis of RNA expanded repeat disorders.

## Results

### Expanded CUGs alter CoQ requirements for normal cellular function

We used the transgenic *C. elegans* strains expressing GFP in the body wall muscle cells with a 3’UTR encoding either 123CTG repeats (123CUG) or none (0CUG)(Garcia et al., 2014) to perform RNAi screens of chromosome 1 to identify modifiers of expanded CUG repeat RNA toxicity (Fig. S1). Of the initial 1824 gene inactivations tested, 8 were identified as altering the defective motility of 123CUG animals with the strongest phenotype due *coq-1* down-regulation. 123CUG animals fed *coq-1* RNAi had a severe paralysis phenotype as 2-day old adults followed by death at 4-days old. In contrast, 0CUG animals had a mild decrease in motility (Fig. 1A, B). In addition, 123CUG animals were egg-laying defective (Egl), exhibiting an accumulation of unlaid eggs in the gonads (Fig. S2B). An independent set of transgenic strains expressing 123CUG and 0CUG repeats displayed the same phenotypes as in our screen (Fig. S2A, B), ruling out the effect of background mutations.

**Figure 1:**
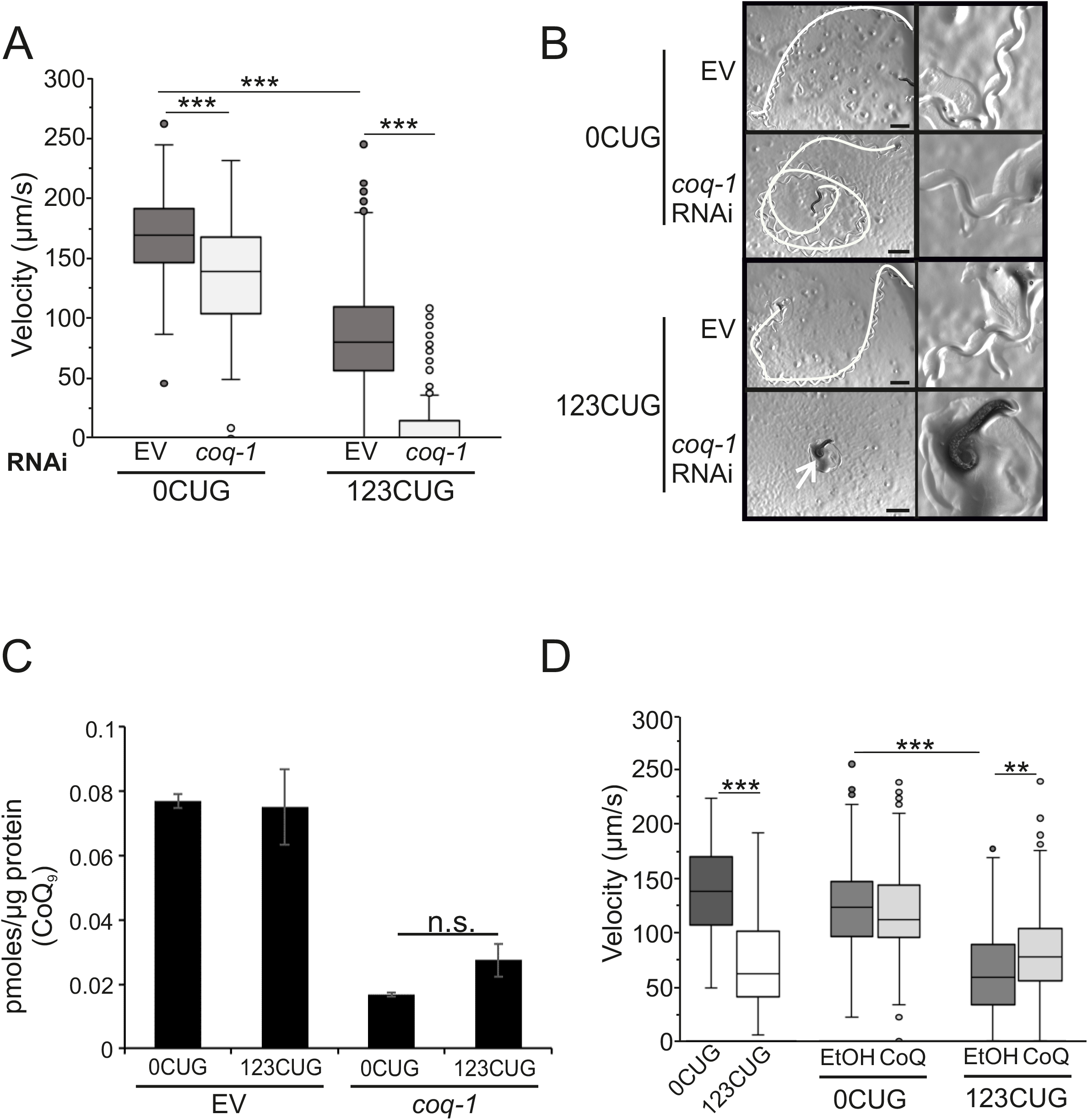
COQ-1 is a suppressor of 123CUG toxicity (A, D) Motility assays; three independent experiments/assay. (A) Velocity of 0CUG and 123CUG’s in empty vector (EV) and *coq-1* RNAi, n≥110 animals/condition. (B) Representative DIC microscopy images of 0CUG and 123CUG’s fed EV or *coq-1* RNAis. White line indicates visible animal tracks, arrow indicates paralyzed animal. Scale bar, 1mm. (C) CoQ_9_ lipid levels of 0CUG and 123CUG fed EV or *coq-1* RNAi. Bars correspond to the mean ± S.E.M. *P* value determined by two-tailed Student’s *t*-test. (D) Velocity of 123CUG and 0CUG’s supplemented with 150μg/mL of CoQ_10_, n>100 animals/condition. Statistics (A, D) determined by Wilcoxon statistical test, and Bonferroni correction; **p<0.005, ***p<0.0001. Box: 25^th^ to 75^th^ percentile; whiskers: 1.5* interquartile range; line in the box: median; dots: outliers.

This supports COQ-1 as a suppressor of expanded CUG toxicity. The gene *coq-1* encodes polyprenyl synthetase which catalyzes the first step in Coenzyme Q (CoQ) pathway, synthesizing the lipophilic polyisoprenoid tail and determining its length(Tran and Clarke, 2007). CoQ is most abundant in mitochondrial membranes where it is best characterized for its role as an essential co-factor in the mitochondrial respiratory chain(Mitchell, 1975). Severe down-regulation of *coq-1* decreases CoQ levels and is characterized by paralysis(Rodriguez-Aguilera et al., 2003). Therefore, we considered whether CoQ biosynthesis was impaired by the expression of the 123CUGs. We analyzed two independent strains of 123CUG animals and 0CUG controls for the expression levels of genes in the CoQ biosynthesis pathway and found no significant changes in two days old adults (Fig. S3A, B). Intracellular synthesis is the major source of CoQ, however contributions from the diet become significant in deficiency conditions(Fernandez-Ayala et al., 2005). Next, we investigated CoQ levels directly in our transgenic *C. elegans* models fed either HT115 *E. coli* (Fig. 1C, S3C) or the CoQ-deficient bacteria GD1 (Fig. S3D). Interestingly, 123CUG animals showed no difference in CoQ levels compared to controls (Fig. S3C, D). Whereas down-regulation of *coq-1* by RNAi induced paralysis in 2-day old adults 123CUG animals and a strong reduction in CoQ levels (Fig. 1C). These results suggest that the 123CUG toxicity was not due to the inability to synthesize CoQ.

Next, we considered whether higher CoQ levels could rescue the 123CUG phenotypes. In *C. elegans*, CoQ content is also temperature-dependent with adult wt (wild type) animals grown at 25°C exhibiting twice the CoQ levels than animals grown at 16.5°C(Jonassen et al., 2002). We observed a temperature dependence in the paralysis and Egl phenotypes of the 123CUG animals (Fig. S4A, B). In addition, at 15°C 123CUGs showed an earlier onset of the severe motility phenotype. Identical results were obtained using independent 123CUG and 0CUG strains, ruling out the contribution of strain-specific background mutations to these phenotypes (Fig. S4A, B). Changing CoQ levels by shifting temperatures has the caveat of affecting the expression of unrelated genes(Conti, 2008). Therefore, we tested directly whether CoQ supplementation of 123CUG animals could rescue the motility defect. CoQ isoforms are species-specific, with a isoprenoide side chain of variable length but an unchanged aromatic ring. *C. elegans* synthesize a CoQ_9_ isoform and acquire CoQ_8_ from dietary *E. coli*(Dutton, 2000). In our assays, we used CoQ_10_, which is the predominant isoform in humans with 10 isoprene units and can rescue defects in this biosynthetic pathway(Takahashi et al., 2012) in *C. elegans*. CoQ_10_ is solubilized in ethanol (EtOH) but this solvent on its own mediates deleterious effects in nematodes and may account for the inability to achieve an even stronger rescue. 123CUG animals supplemented with CoQ_10_ showed a robust rescue of the motility defect (Fig. 1D). Although 123CUG animals are not CoQ deficient, the sensitivity of these animals to CoQ levels suggests a new metabolic mechanism to RNA repeat toxicity.

### Expanded CUG RNA repeats disrupt mitochondrial respiration

CoQ is best known for its essential role as a redox carrier(Crane and Navas, 1997) in the electron transport chain (ETC) shuttling electrons from complexes I and II to complex III during oxidative phosphorylation (Fig. S5A). We asked whether a deficit in complexes I and II could modulate the RNA repeat toxicity. Severe reductions in the mitochondrial ETC components can exert deleterious organismal effects(Rea et al., 2007), resulting in small and sick animals for most of the genes tested (*gas-1, nuo-6*, and *mev-1*). To circumvent this problem, we diluted the RNAis to concentrations low enough so as not to affect animal size or result in lethal, sick phenotypes but yet still induced a toxicity effect, such as motility defect (Fig. S5B). In contrast, gene inactivation in the 123CUGs did not result in further motility defect, suggesting the presence of impaired ETC components.

Next, we addressed whether mitochondrial respiration was affected. We measured mitochondrial oxygen consumption rates (OCR) (Fig. S5C) of 2-day old adult nematodes and found that the 123CUG animals showed increased basal respiration, which reduced the spare respiratory capacity (Fig. 2A, B). These results are indicative of uncoupled respiration and dissipation of the membrane potential. Thus, the reduced respiratory capacity in the 123CUGs decreased the ability of these animals to respond to bioenergetic demands. To test whether inactivation of the CoQ pathway would further enhance 123CUG’s respiration deficiency, we treated our strains with *coq-1* RNAi that generated a severe decline in basal and maximal respiration consistent with a collapse of the ETC function (Fig. S5D). We reasoned that if CoQ impairment is a key modulator of respiration dysfunction in the 123CUG strains then supplementation might reverse the phenotype as was observed with impaired motility (Fig. 1D). Indeed, CoQ supplementation to the 123CUG animals reduced basal respiration (Fig. 2C). Together, these findings indicate that expression of the 123CUG induces mitochondrial uncoupling that can be rescued with CoQ supplementation.

**Figure 2:**
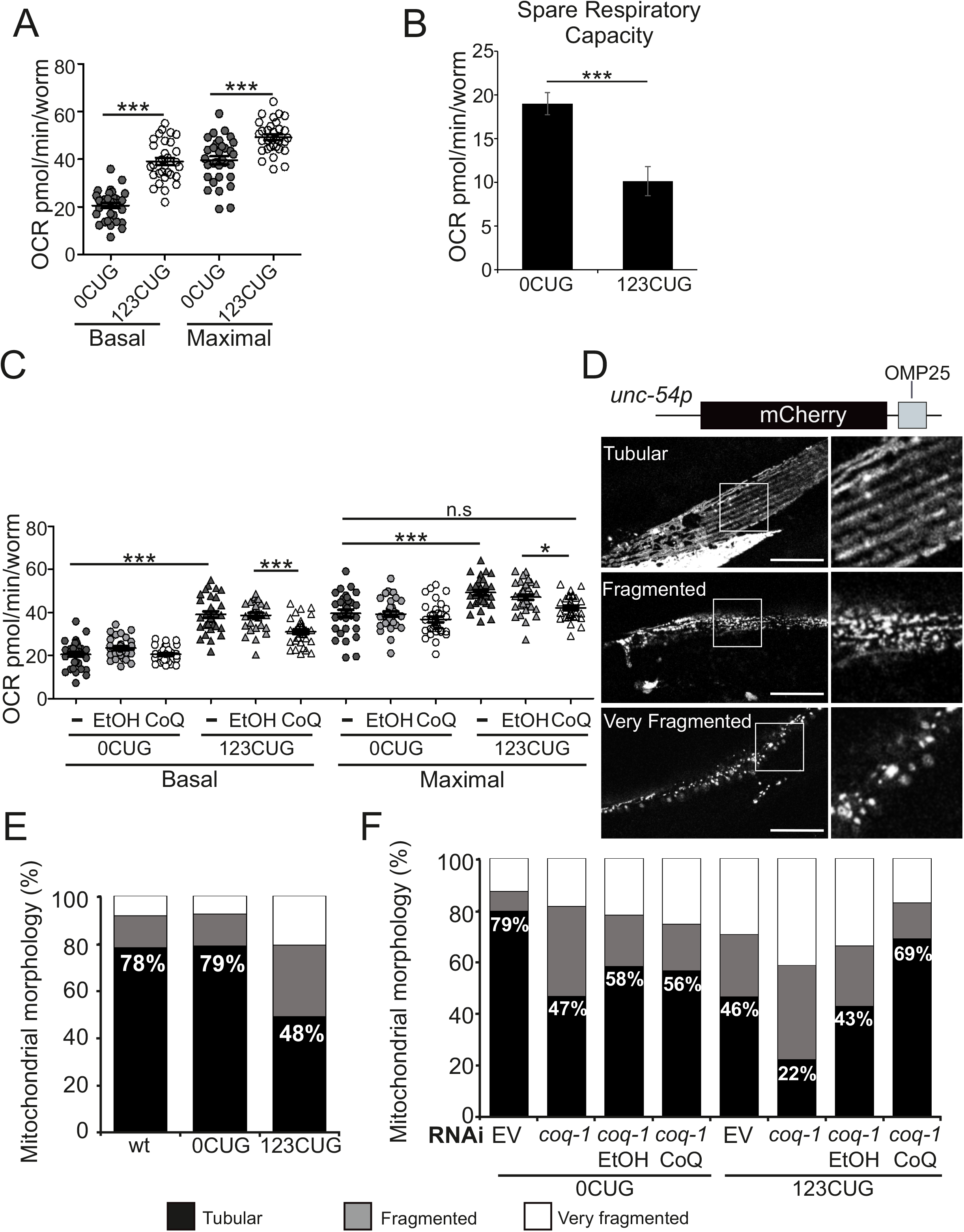
Expanded repeats cause mitochondrial dysfunction (A, B) Mitochondrial respiration of 123CUG and 0CUG; three independent experiments/assay. The uncoupler FCCP was used to measure maximal respiration. Bars correspond to mean ± S.E.M. (A) 123CUG and 0CUG animals’ without any treatment. A two-tailed Student’s t-test was used with Bonferroni correction. ***p<0.0001, n=120 animals/experiment. (B) Spare respiratory capacity of 123CUG and 0CUG. Bars represent mean ± S.E.M. Two-tailed Student’s t-test was used to analyze statistical significance; ***p<0.0001. (C) Effect of CoQ_10_supplementation on respiration of 0CUG and 123CUG; 75μg/mL CoQ_10_,n≥100 animals/experiment. Statistical significance determined by two-way Anova with Bonferroni correction; **p<0.005, ***p<0.0001. (D) Schematic representation of mitochondrial reporter construct and confocal images of the different mitochondrial morphologies observed. Scale bar, 25μm. (E–F) Quantification of mitochondrial morphology of 2-days old adult 123CUG and 0CUG. (D) Without any treatment; five independent experiments, n≥96 mitochondria/condition. (E) *coq-1* downregulation and supplementation with 75μg/mL of CoQ_10_. Two independent experiments performed, n≥39 mitochondria/condition.

### 123CUG causes altered mitochondrial morphology

There is a close interplay between the mitochondrial respiratory chain function and membrane dynamics(Youle and van der Bliek, 2012). Most notably, mitochondrial uncoupling is associated with dramatic remodeling of organelle morphology into a fragmented state. To test whether mitochondrial morphology was affected by expression of the 123CUG repeats, we generated a body-wall muscle-specific mitochondrial fluorescent reporter expressing mCherry with the OMP25 outer-membrane targeting sequence (Fig. 2D). The reporter was crossed into both 123CUG and 0CUG strains, and mitochondrial morphology investigated in 2-day old adults. We classified the morphologies into three categories: tubular, fragmented, and very fragmented (tubules were no longer visible) (Fig. 2D). 123CUG animals showed a dramatic decrease in tubular mitochondria (Fig. 2E). In the muscle, mitochondrial morphology has been associated with the copy number alterations of mitochondrial DNA (mtDNA)(Chen et al., 2010). However, we found that increased mitochondrial fragmentation did not affect mtDNA levels (Fig. S5E). Next, we tested how CoQ modulation affected the mitochondrial morphology of 123CUG and 0CUG animals. Consistent with the previous phenotypes, modulating CoQ levels in animals expressing 123CUG repeats improved or exacerbated the mitochondrial morphology defect (Fig. 2F). Together, the data show that the alteration in mitochondrial morphology correlates with mitochondrial dysfunction and can be rescued by CoQ supplementation.

To establish whether the RNA repeat toxicity on mitochondrial morphology is a conserved cellular mechanism, we investigated the effects of expressing GFP with a 3’UTR bearing 0, 5, 100 or 200CTG repeats in the human HeLa cell line (Fig. 3A). Cells were analyzed 20 hours post-transfection of the constructs and mitochondrial morphologies were classified into four categories: tubular, intermediate, fragmented or aggregated (Fig. 3B). We observed that expression of the expanded repeats altered mitochondrial morphology, progressively reducing the percentage of tubular mitochondria as the length of the repeat increased (Fig. 3C). Taken together, these results support a conserved role of RNA repeat toxicity on mitochondrial dysfunction.

**Figure 3:**
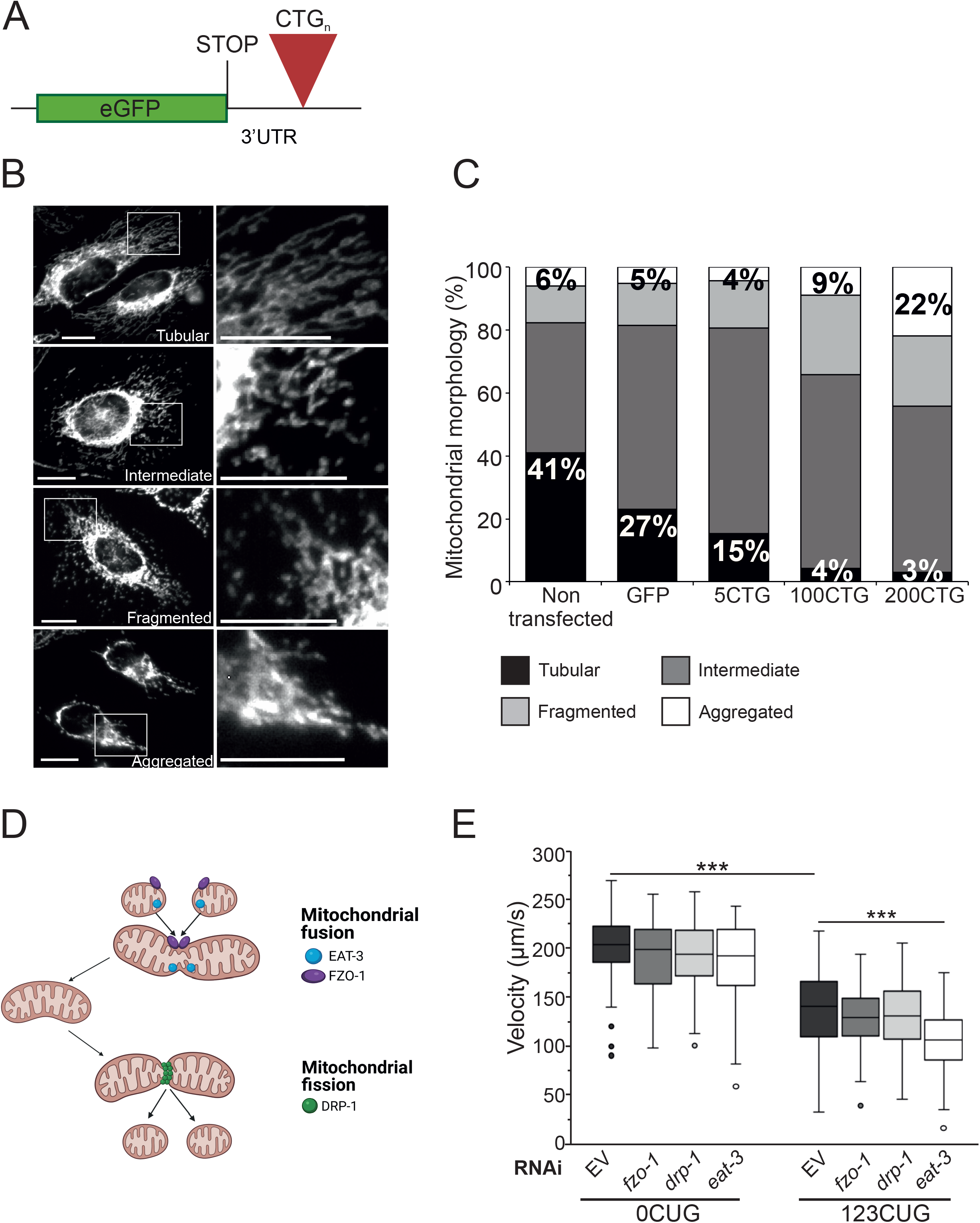
Expanded repeats disrupt mitochondrial morphology. (A – C) Mammalian mitochondrial morphology is dependent on repeat length. (A) Schematic representation of GFP construct bearing CTG repeats in 3’UTR for mammalian expression. (B) Representative microscopy images of the mitochondrial morphologies visualized. Scale bar, 20μm. (C) Quantification of mitochondrial morphologies of GFP and GFP bearing 5CTG, 100CTG and 200CTG repeats. (D – E) Mitochondrial dynamics is altered in 123CUG animals.(D)Schematic representation of mitochondrial dynamics (Created with BioRender.com). A steady-state of mitochondrial dynamics is achieved by a balance between fusion and fission. During fission, DRP-1 is recruited from the cytosol and binds to the mitochondrial outer membrane constricting it and leading to the separation of the mitochondria into two. Mitochondrial fusion requires both inner and outer membranes from individual mitochondria to fuse. This process is regulated by EAT-3 and FZO-1, respectively. (E) Velocity of 0CUG and 123CUG strains upon inactivation of mitochondrial dynamics regulatory genes. Statistical significance was determined by Wilcoxon statistical test, with Bonferroni correction; ***p<0.0001. Two independent experiments were performed, n≥90 animals/ condition. Box: 25^th^ to 75^th^ percentile; whiskers: 1.5 * interquartile range; line in the box: median; dots: outliers.

Because mitochondria dynamics is dependent on the balance between fusion and fission (van der Bliek et al., 2013; Yu et al., 2020), we asked whether these pathways modulated mitochondrial dysfunction in 123CUG animals. To address this, we used RNAi to silence the factors required for mitochondrial fission, (*drp-1*) and fusion (*fzo-1* and *eat-3*) in 123CUG and 0CUG strains (Fig. 3D). No significant changes in motility were detected in the 123CUG or 0CUG worms upon inactivation of *fzo-1* or *drp-1*, which regulate the fusion and fission of the mitochondrial outer membrane, respectively (Fig. 3E). In contrast, we found that *eat-3* RNAi further impaired the motility defect in the 123CUG animals. EAT-3 regulates the inner mitochondrial membrane dynamics and is known to be sensitive to uncoupled respiration(Kanazawa et al., 2008).

### The RNA-binding factor MBL-1 mediates mitochondrial disruption

In myotonic dystrophy disorders the loss of the trans-acting MBNL RNA-binding factors is proposed to be a major contributor of pathogenesis(Miller et al., 2000). Since MBNLs/MBL-1 regulate the splicing of hundreds of transcripts(Du et al., 2010; Norris et al., 2017; Wang et al., 2012), a model has been proposed whereby depletion of these RNA-binding proteins dysregulates RNA splicing and contributes to the molecular pathogenesis. Thus, we addressed whether MBL-1 was connected to the metabolic disruption in mitochondrial function with expression of the 123CUG repeats. For these experiments, we used a *mbl-1* null allele mutant strain to test for effects on motility. The *mbl-1* mutants exhibited a strong decrease in velocity as 2-day old adults (Fig. 4A). The *mbl-1(tm1563)* animals also showed the same temperature dependence on motility, Egl, CoQ requirement and unaltered CoQ levels that we observed with the 123CUG animals (Fig. 4A and S6A, B, C). Together, the data strongly suggest the interplay between CoQ metabolism and MBL-1.

**Figure 4:**
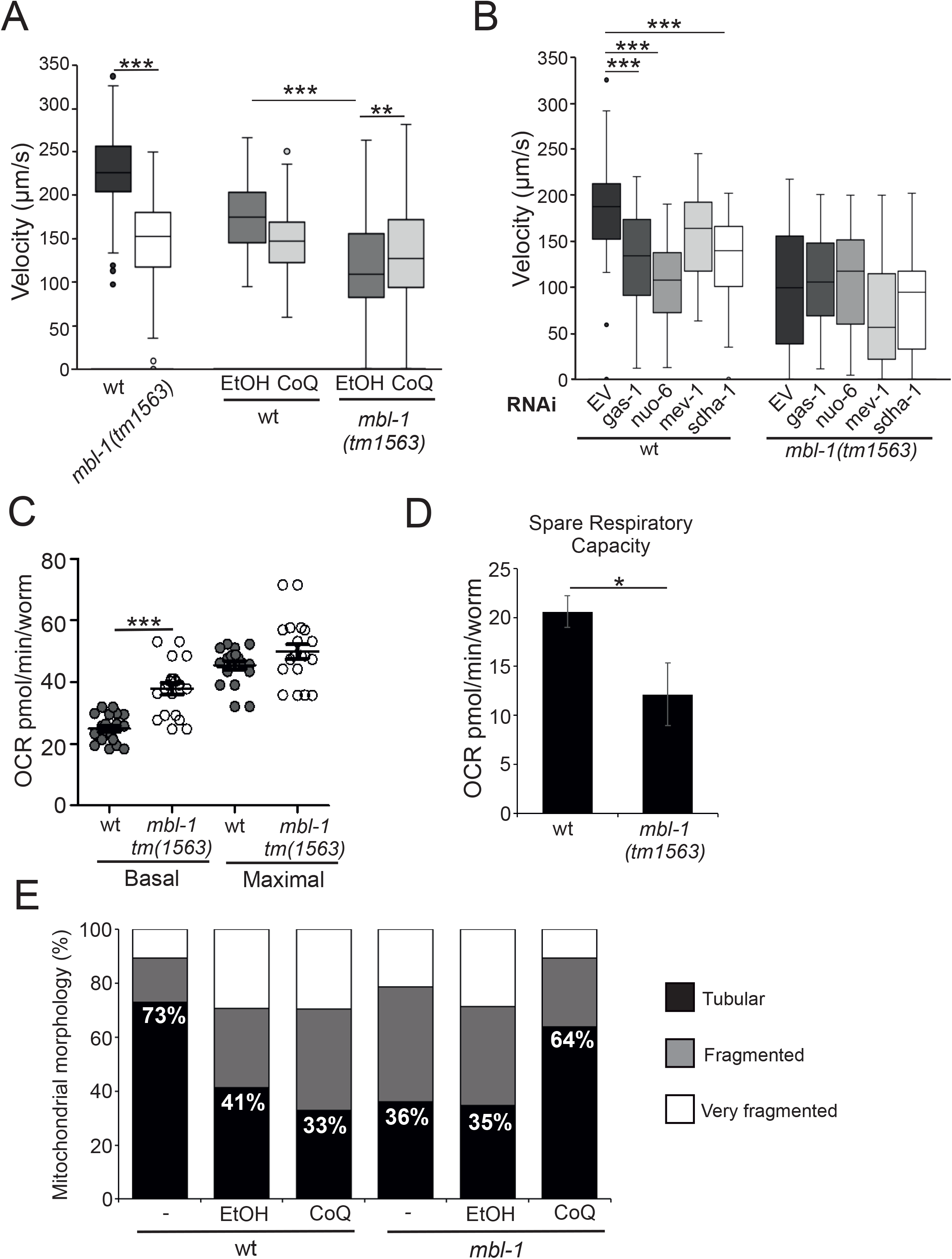
Mitochondrial dysfunction is dependent on MBL-1. (A-B) Motility assays of wt and *mbl-1(tm1561)* strains. *P* value determined by Wilcoxon statistical test, with Bonferroni correction. **p<0.005, ***p<0.0001. Box: 25^th^ to 75^th^ percentile; whiskers: 1.5* interquartile range; line in the box: median; dots: outliers. (A) Velocity of animals fed EV and supplemented with 75μg/mL CoQ_10_. Two independent assays performed, n≥94 animals/condition. (B) Velocity of wt and *mbl-1(tm1563)* fed RNAi of *gas-1, nuo-6, mev-1* and *sdha-1*. RNAi clones were diluted with EV to the final concentrations: 20% (*gas-1, nuo-6*) and 40% (*mev-1*). n≥44 animals/condition. (C) Respiration assay of wt and *mbl-1(tm1563)* strains. Two independent experiments performed, n=120 animals/experiment. Bars correspond to mean ± S.E.M. Statistical significance determined by two-tailed Student’s *t*-test with Bonferroni correction. (D) Spare respiratory capacity of wt and *mbl-1(tm1563)*. Bars represent mean ± S.E.M. Two-tailed Student’s t-test was used to analyze statistical significance; *p<0.05 (D)Quantification of mitochondrial morphology of wt and *mbl-1(tm1563)* supplemented with 75μg/mL CoQ_10_. At least three independent experiments performed, n>105 mitochondria/condition.

Next, we tested *mbl-1* mutants for impairment of ETC complexes I and II. Down-regulation of *gas-1, nuo-6* (complex I), and *sdha-1* (complex II) had no effect on the *mbl-1* mutant strain (Fig. 4B), indicating that these components are already compromised on *mbl-1(tm1563)* animals and further supporting a link between MBL-1 and ETC function. We monitored the rates of oxygen consumption of 2-day old adults and found that *mbl-1* mutants had an increased basal respiration and reduced spare respiratory capacity (Fig. 4C,D). Together, these results showed that *mbl-1(tm1563)* mutants have dysfunctional OXPHOS.

To test whether *mbl-1* mutants also showed changes in mitochondrial morphology, we crossed the mitochondrial reporter (Fig. 2D) to the *mbl-1(tm1563)* strain. In 2-day old adults, *mbl-1(tm1563)* animals displayed a reduction in tubular mitochondria, which we could rescue with CoQ supplementation (Fig. 4E). Importantly, CoQ supplementation rescues the tubular phenotype, a morphology close to wt animals grown on media without solvent solution. Also, we found that *mbl-1(tm1563)*’s increased mitochondrial fragmentation did not affect mtDNA levels (Fig. S6D). Together these data reveal that MBL-1 disruption contributes to mitochondrial dysfunction and the membrane morphology defect.

### RNA repeat expansions engage the Ribosome Quality Control complex

Since the import of most of the mitochondrial proteome requires a membrane potential, uncoupled respiration will severely impair the process and generate a proteotoxicity on the organelle surface. Consistent with this principle, it has been reported that ectopic induction of mitochondrial dysfunction triggers translocation of factors in the ribosome-associated quality control (RQC) pathway to the organelle(Heo et al., 2010). Our data led us to consider whether the RQC surveillance system plays a role in the modulation of 123CUG RNA repeat phenotypes. To investigate this hypothesis, we tested whether components of the RQC pathway modify the expanded CUG toxicity. Vms1 is a release factor that translocates to mitochondria in response to a variety of organelle stressors(Izawa et al., 2017) and interacts with cdc48, an AAA ATPase, which delivers ubiquitylated polypeptides to the proteasome for degradation. Inactivation of *vms-1* and *cdc-48*.*2* in 123CUG animals considerably worsened the motility defect, inducing a severe paralysis in the animals fed *cdc-48*.*2* RNAi (Fig. 5A). In contrast, the effect on motility in the 0CUG animals was mild. To test whether *mbl-1* mutants would exhibit a similar requirement, we down-regulated *cdc-48*.*2* in *mbl-1(tm1563)* animals because it had the strongest effect on motility. Inactivation of *cdc-48*.*2* resulted in a strong inhibition of *mbl-1* mutant motility (Fig. 5B). Together, our findings reveal that expression of the 123CUG RNA repeat induces and triggers a proteotoxicity on the organelle surface that requires the RQC to resolve.

**Figure 5:**
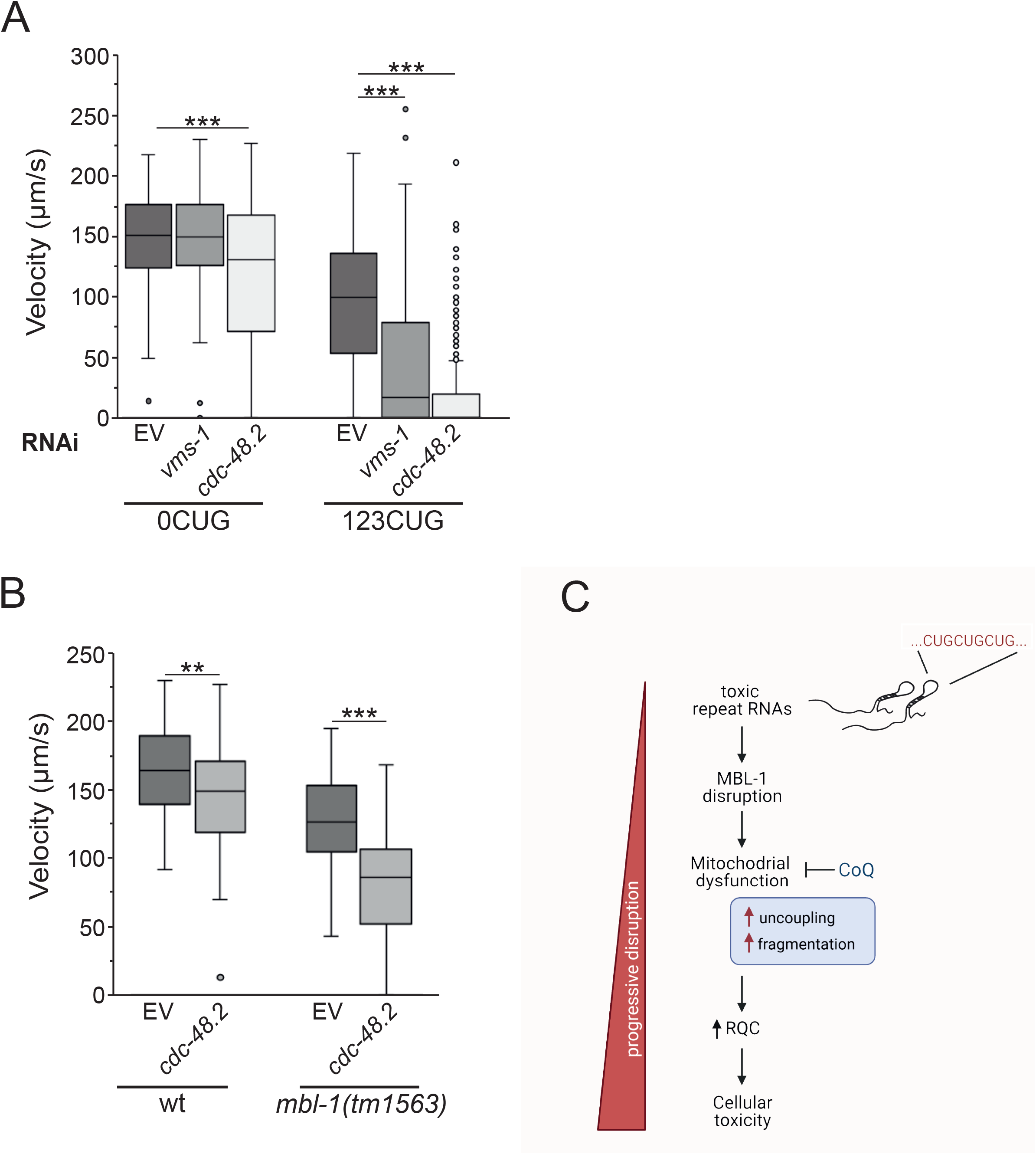
RQC function is required in 123CUG animals to counteract repeat toxicity. (A-B) Motility assays, two independent experiments performed; *P* value determined by Wilcoxon statistical test, with Bonferroni correction. **p<0.005, ***p<0.0001. Box: 25^th^ to 75^th^ percentile; whiskers: 1.5* interquartile range; line in the box: median; dots: outliers. (A) 0CUG and 123CUG fed *vms-1* and *cdc-48*.*2* RNAis. (B). wt and *mbl-1(tm1563)* fed *cdc-48*.*2* RNAi. Model of mitochondrial dysfunction by toxic repeats. Expression of transcripts bearing expanded CUG repeats leads to MBL-1 association and mitochondrial dysfunction, with disruption of respiration and increased organelle fragmentation. The progressive accumulation of dysfunctional mitochondria trigger the recruitment of the RQC quality control complex and result in cellular toxicity.

## Discussion

Our genetic screen for modifiers of RNA repeat toxicity reveals a new pathogenic mechanism. Here we report that expression of expanded CUGs repeats in the body wall muscle cells induces a motility defect caused by mitochondrial dysfunction. In turn, this organelle stress triggers a cytosolic proteotoxicity that requires the RQC to resolve. Moreover, our study shows how MBL-1, a *bona fide* factor in RNA repeat toxicity, is linked to these two phenotypes, with mitochondrial dysfunction dependent on MBL-1. Together, our findings demonstrate a novel, conserved link between toxic RNAs, mitochondrial dysfunction and the quality control mechanisms required for cell homeostasis (Fig. 5C).

Our data show that the mitochondrial dysfunction is a primary toxic event and is surprisingly not due to reduced CoQ levels. In *C. elegans*, mitochondrial biogenesis in body wall muscle cells occurs during embryogenesis(Cornaglia et al., 2015), during the transition from the third to fourth-larval stages(Tsang and Lemire, 2002). In our 123CUG animals, RNA expression under the *myo-3* promoter also starts during embryonic development, when transcripts likely start to accumulate. However, the mitochondrial dysfunction we observe does not appear to be mediated via impairment in organelle biogenesis, as evidenced by the CoQ levels and mtDNA copy number of both 123CUG and *mbl-1* mutant animals. Instead, we favor the following model. Since RNA repeat toxicity is characterized by progressive degeneration, which we observe with the age-dependent deterioration of the 123CUG’s motility(Garcia et al., 2014), we hypothesize that there is a threshold for the accumulation of the expanded transcripts before inducing progressive mitochondrial dysfunction.

Mitochondria participate in numerous metabolic pathways, and changes in bioenergetic efficiency are well-known to affect mitochondrial dynamics and protein import(Liesa and Shirihai, 2013). Because of the strict demand to import over 1000 proteins synthesized on cytosolic ribosomes into mitochondria, failures in the bioenergetic status of the organelle can exert profound effects on cytosolic protein homeostasis and the requirement for RQC. A central function of RQC is the quality control of protein synthesis, contributing to the elimination of aberrant mRNAs and to promote nascent chains ribosome recycling(Joazeiro, 2019). Therefore, intracellular rerouting of these factors due to mitochondrial dysfunction will likely have a deleterious and cumulative effect on cytosolic protein homeostasis. Furthermore, we predict that as an organism ages, the reliance on the RQC surveillance system for cellular proteostasis increases, particularly as transcription errors are known to accumulate, leading to a compromised RQC surveillance.

Based on our findings, we propose that mitochondrial dysfunction arising in RNA repeat-based disorders may be a reoccurring mechanism that deserves a more thorough consideration going forward. Our rationale is the following. Although, a role for mitochondrial involvement in the pathogenesis of DM1 patients has previously been made based upon reduced enzymatic activity these are often associated with conflicting results (Gramegna et al., 2018; Vita et al., 1993). Nonetheless, mitochondrial abnormalities including “ragged fibers” with enlarged mitochondria and impaired muscle oxidative metabolism have been observed in biopsies from myotonic dystrophy (DM) patients(Gramegna et al., 2018; Ono et al., 1986). Thus, we suggest that mitochondrial dysfunction may represent a more significant contributor to RNA-repeat toxicity than previously thought. Similarly, MBNL has been in implicated in an increasing number of repeat-based disorders, from DM to Huntington disease-like 2 (HDL2)(Rudnicki et al., 2007). Our data showed that MBL-1 disruption affects mitochondrial function. Moreover, MBNL/MBL-1 sequestration rates are known to correlate with RNA repeat lengths(Thomas et al., 2017), which is supported by the fragmentation phenotypes we observed (Fig. 3C). Together these data provide new insights to the molecular roles played by MBNL/MBL-1 in repeat-based cellular toxicity.

Our data now underscores a consistent molecular mechanism by which mitochondrial dysfunction is a central contributor to pathogenesis. Finally, because our results show that these pathogenic processes can be ameliorated, with a significant improvement in the toxic effects and on organismal fitness, similar pharmacological approaches targeting the mitochondria may represent viable future therapeutic options.

## Methods

### *C. elegans* Strains

*C. elegans* were grown following standard culture protocols(Brenner, 1974), at 20°C unless otherwise indicated. The following strains were used: N2 Bristol as the wild-type strain, *mbl-1(tm1563)* (FX01563), strains expressing GFP with 123CUG repeats in their 3’UTR(Garcia et al., 2014) (GR2024, GR3208) or GFP alone (GR2025, GR3207)(Garcia et al., 2014). A mitochondrial reporter strain was generated expressing the fluorophore mCherry with a mitochondrial targeting sequence (GAR118). GAR118 was generated by injecting the mCherry transgene into wt animals at the concentration of 50ng/μL.

### Plasmids and constructs

The mitochondrial outer membrane protein-25 (OMP25) sequence(Nemoto and De Camilli, 1999) was cloned by PCR amplification into the *C. elegans* pPD49.26 plasmid, bearing an *unc-54* body wall muscle specific promoter, together with a mCherry fluorophore. The PCR primers used to clone the OMP25 into the pPD49.26 plasmid are in Supplementary Table 2.

### RNA Interference (RNAi)

RNAi bacteria expressing dsRNA were grown overnight in LB media and induced the following day by the addition of 500μL/L IPTG before cultures were plated, as previously described(Timmons et al., 2001). For *coq-1* RNAi, a higher IPTG concentration of 6mL/L was used. Strains to be analyzed by RNAi were synchronized by hypochlorite (NaOCl) treatment of gravid adults, their eggs collected, plated onto the RNAi plates and left to develop. At the L4 stage, animals were transferred onto new RNAi plates containing 10μM of 5-fluorouracil. RNAi dilutions were performed by measuring the bacterial optical density (OD_600_) and RNAi was diluted with EV. The RNAi dilutions were calculated in a way that the RNAi concentration was low enough not to affect the animals development, but high enough to show toxic phenotypes.

### RNAi Screens

RNAi-mediated gene inactivation was achieved through the feeding method(Kamath et al., 2001). Animals were synchronized by NaOCl bleaching and overnight hatching in M9. Briefly, in the RNA degradation screen RNAi clones were grown overnight in 96-well 2mL blocks containing LB with 200μg/mL carbinicillin and then induced with 4mM of isopropylthiogalactoside (IPTG) for 3h. The RNAi cultures were pelleted and then dissolved to a 9x concentration in S-basal medium, and 10μl of RNAi bacteria was transferred to each well in a 96-well plate. The synchronized L1s were filtered and aliquoted (30μl) into the RNAi-containing 96-well plates (16 animals/well) and allowed to develop to the L3-stage larvae at 20°C. We screened the first two-thirds of chromosome I by RNAi, corresponding to 1824 genes, using a reporter strain expressing a GFP with a GC-rich sequence in its 3’UTR, whose RNA is targeted for degradation(Garcia et al., 2014). Animals were screened for the presence of fluorescence and scored 0 to 3 based on the intensity of the GFP signal detected, with 0 having no GFP signal and 3 the strongest signal (Fig. S1). Animals fed the empty vector L4440 or targeting *smg-2* were used as negative and positive controls, respectively. All positives were re-screened twice, and inactivations with a score above 1.5 and a human orthologue were used for a motility screen, 32 RNAis. The motility screen was performed as described (Fig. S1), the synchronized L1s were transferred onto 10cm RNAi plates where animals were allowed to develop to 2 days adults (5 days after synchronization). Plates containing L4440 or *smg-2* RNAi bacterial clones were used as controls. Animals were analyzed for changes in locomotion, with the clones re-screened in triplicate. After re-screening, 8 RNAis were identified as altering the defective motility of 123CUG animals, without similarly affecting the 0CUGs control. The RNAi clones were verified by sequencing the inserts.

### *C. elegans* motility assays

For the motility screen, gravid adults were synchronized by NaOCl treatment, and hatched overnight in M9. Around 150 L1 larval-stage animals were plated. Since overnight hatching implies animal starvation and it is known that CoQ levels decrease with dietary restriction, we started plating the eggs after the synchronization. Approximately 200 eggs were plated onto seeded plates. 2-day old adults animals were collected and rinsed three times with M9 buffer, transferred onto 10cm agar plates without bacteria, and let rest for 30min prior to imaging. For each assay a minimum of seven 25s videos were recorded using an Axio Zoom.V16 microscope, at 8x magnification. The data was analyzed to determine the animals’ speed (μm/s) using the Matlab program developed by M. Crivaro at the Light Microscopy Unit (Institute of Biotechnology, University of Helsinki).

### Fluorescent and DIC Imaging

All *C. elegans* images were of 2-day old adults animals (4 days post-hatching) unless indicated otherwise. *C. elegans* motility images were taken using an Axio Zoom.V16 microscope, at 20x magnification. Animals were picked onto plates with 10x concentrated OP50 and imaged 2min post-transfer. Images of Egl phenotype were taken using a Zeiss Imager.M2 microscope, at 10x objective. Nematodes to be assayed were transferred onto a 2% agarose pad, on a glass slide, and immobilized with 10mM of levamisole.

Mitochondrial morphology images were obtained by mounting, and immobilizing the animals expressing the mitochondrial reporter with 10mM levamisole on a 5% agarose pad. Imaging was performed using a Leica SP8 confocal microscope, with 93x objective. The images were analyzed in a blind manner for their morphology when classified in the different categories.

### Mitochondrial respiration assays

Oxygen consumption rates (OCR) were measured using a Seahorse XFe96 Analyzer. Gravid adults were synchronized, the eggs collected and plated on RNAi plates seeded with EV or *coq-1* RNAis, and OCR measured when animals reached 2-day adulthoods. The Seahorse XF Sensor Cartridges were hydrated with XF Calibrant Solution and these were maintained over-night at 27°C in a non-CO_2_ incubator(Koopman et al., 2016). Prior to the assay, the XF Sensor Cartridge was calibrated. The animals were picked into individual wells of a Seahorse 96-well XF Cell Culture microplate, 10 worms/well in 80μL of M9 buffer, and a minimum of at least 10 wells per condition was used. OCR measuring cycles consisted of 3 min mixing, 30sec of rest, and 3 min of analyzing oxygen consumption. Four measurement cycles were performed for each condition: basal OCR, OCR upon FCCP treatment and OCR upon sodium azide treatment. Animal number, FCCP and sodium azide concentrations were optimized for this study.

### CoQ supplementation assays

In CoQ supplementation assays animals were fed EV or the CoQ-deficient GD1 strain of *E. coli* bacteria. In assays with GD1 bacteria, animals were grown on GD1 for at least two generations prior to the assay to eliminate possible confounding effects from dietary CoQ. CoQ supplementation was performed by dissolving the CoQ_10_ in ethanol and adding it to the agar at the desired concentration. For motility assays, animals were then synchronized and the eggs transferred onto plates seeded with GD1 and containing 150μg/mL of CoQ_10_ (Sigma). Motility analysis was performed as described. Alternatively, motility assays were performed on plates seeded with EV and supplemented with 150μg/mL or 75μg/mL of CoQ_10_. The effect of CoQ supplementation on respiration and on mitochondrial morphology was examined by feeding the animals EV or *coq-1* RNAi and supplementing the plates with 75μg/mL of CoQ_10_. Imaging and respiration analysis were performed as described.

### Quantitative RT-PCR (qRT-PCR)

Animals were collected from three biological replicas, as 2-day old adults, and frozen in liquid nitrogen. For total RNA extraction, TRIzol Reagent (Ambion) was used followed by chloroform extraction and isopropanol precipitation. cDNA was synthesized from 250ng RNA using QuantiTect Reverse Transcription Kit (Qiagen), and qRT-PCR reactions were performed with SYBR Green reagent (Roche) using Lightcycler 480 (Roche). qRT-PCR data was normalized to *cdc-42* and *pmp-3* gene expressions. The 2^-ΔΔct^ method was used for comparing relative levels of mRNAs. Primers are listed in Supplementary Table 2.

### Mitochondrial copy number

Mitochondrial copy number was determined by qPCR of 2-day adult samples, as described earlier(Bratic et al., 2010). Briefly, animals were collected in lysis buffer (50mM KCl, 10mM Tris-HCL ph 8.3, 2.5mM MgCl_2_, 0.45% NP-40, 0.45% Tween-20, 0.01% gelatin and 60μg/mL protease K) and incubated at 60°C for 2h followed by 15min at 95°C. qRT-PCR was performed with SYBR Green reagent (Roche) using Lightcycler 480 (Roche) and expression data was normalized to the *act-1* levels. Primers are listed in Supplementary table 2.

### Sample preparation for lipid analysis

Three biological samples containing around 1000 animals each, were collected as 2-day adults and transferred to 2mL polypropylene tubes with 250μL of lipid lysis buffer (20mM Tris-HCl pH 7.4, 100mM NaCl, 0.5mM EDTA, 5% glycerol) and kept on ice for 15 minutes. Sample homogenization was performed using a Precellys 24 homogenizer at 6400rpm, twice for 10sec with 5sec interval. This homogenization procedure was repeated twice and samples were kept on ice between homogenizations.

### Lipid extraction

Lipid extraction was performed as described previously(Pellegrino et al., 2014) with some modifications. To 20μl of the worm lysate, 1ml of a mixture of Methanol: Methyl tertiary-butyl ether: Chloroform (MMC) 1.33:1:1 (v/v/v) was added. The MMC was fortified with the SPLASH mix of internal standards (Avanti Lipids). After brief vortexing, the samples were mixed continuously in a Thermomixer (Eppendorf) at 25°C, for 30min at 950rpm. Samples where then centrifuged for 10min, 16000g, 25°C to obtain protein precipitation. The single-phase supernatant was collected, dried under N_2_, and stored at –20°C until analysis. Before analysis, the dried lipid samples were redissolved in 100μL MeOH:Isoproanol (1:1).

### Liquid chromatography mass spectrometry conditions

Liquid chromatography was done as described previously(Cajka and Fiehn, 2016) with some modifications. Lipids were separated using C18 reverse phase chromatography. Vanquish LC pump (Thermo Scientific) was used with the following mobile phases; A) Acetonitrile:Water (6:4) with 10mM ammonium acetate and 0.1% formic acid and B) Isopropanol: Acetonitrile (9:1) with 10mM ammonium acetate and 0.1% formic acid. The Acquity BEH column (Waters) with the dimensions 100mm*2.1mm*1.7μm (length*internal diameter*particle diameter) was used. The following gradient was used with a flow rate of 0.6 ml/minutes; 0.0-2.0 minutes (isocratic 30%B), 2.0-2.5 minutes (ramp 30-48% B), 2.5-11 minutes (ramp 48-82%B), 11-11.5 minutes (ramp 82-99%), 11.5-12 minutes (isocratic 100%B), 12.0-12.1 minutes (ramp 100-30% B) and 12.1-15 minutes (isocratic 30%B).

The liquid chromatography was coupled to a hybrid quadrupole-orbitrap mass spectrometer (Q-Exactive HFx, Thermo Scientific). A full scan acquisition in negative and positive ESI was used, scanning from 200-2000 m/z at a resolution of 120000 and AGC Target 1e6, max injection time 200ms. Data-dependent scans (top10) were acquired using normalized collision energies (NCE) of 20, 30, 50 and a resolution of 15,000 and AGC target of 1e5.

### Ubiquinone identification

Identification of the quinone lipids was achieved using four criteria; 1) high accuracy and resolution with an accuracy within m/z within 5ppm shift from the predicted mass, and a resolving power 70000 at 200 m/z. 2) Isotopic pattern fitting to expected isotopic distribution. 3) comparison of the retention time to an in-house database and 4) fragmentation pattern correspondence to an in-house experimentally validated lipid fragmentation database. Quantification was done using single point calibration by comparing the area under the peak of different quinones to the area under the peak of SPLASH internal standard closed in time to the area under the peak of the internal standard and then normalized to protein concentration. Mass spectrometric data analysis was performed in Tracefinder software 4.1 (ThermoScientific) for peak picking, annotation and matching to the in-house fragmentation database.

### Mammalian Cell Culture

HeLa cells were grown at 37°C and 5%CO_2_ in Dulbecco’s Modified Eagles Medium (DMEM). Media was supplemented with 10% fetal bovine serum, 1× glutamax and 50 μg/ml uridine.

### Immunoblotting and antibodies

HeLa cells were grown on coverslips and transfected, upon 50-60% confluence, with 2μg of GFP, 5CTG, 100CTG or 200CTG using JetPrime. The media was changed 4h post-transfection, and 20h post-transfection the cells were rinsed three times in phosphate buffered saline (PBS) solution and then fixed in 4% paraformaldehyde for 15 min. After fixation, cells were washed in PBS and incubated with 100% cold methanol for 5 min and washed again in PBS. Cells were then blocked in 5% bovine serum albumin (BSA)/PBST (PBS + 0.1% Tween 20) and incubated with mouse anti-sdha antibody (C2061 MitoSciences/Abcam) 1:250 for 1h, at room temperature. Cells were washed three times in PBST and incubated with anti-mouse Alexa 594 1:1000 (Life Technologies) for 1h, at room temperature. Cells were washed three times in PBST and mounted with DABCO/MOWIOL on glass slides for imaging. A Zeiss Imager.M2 microscope was used for imaging with a 40x objective.

### Statistical Analysis

Statistical analysis of motility assays was performed using the Wilcoxon statistical test with the Bonferroni correction. A minimum of two independent experiments were performed per gene and condition tested. Statistical analysis of mitochondrial respiration was performed using the two-tailed Student’s t-test or the two-way Anova statistical test if different conditions or genes were analyzed. The Bonferroni correction was used with both statistical tests. A minimum of two independent experiments were performed in mitochondrial respiration assays, per gene or condition tested. Statistical significance of the spare respiratory capacity and CoQ_9_ lipid levels was determined using two-tailed Student’s t-test with Bonferroni correction. Statistical analyses for qRT-PCR data were carried out in Excel, and the data represent the mean of 3 biological replicates ± SD. Statistical details can be found in the figures and figure legends. In addition, this statistical data is listed in Supplementary Table 5.

## Data availability

Strains and plasmids are available upon request.

## Acknowledgments

We thank C. F. Clarke (University of California) for providing the GD1 *E. coli* strain, M. Frilander and C. Holmberg (University of Helsinki) for use of tissue culture and incubators, respectively. We are grateful to M. Crivaro, from the Light Microscopy Unit at University of Helsinki, for developing the program used to analyze the *C. elegans* motility data. We thank M. Mahadevan (University of Virginia) for the plasmids bearing the CUG repeats. We are also grateful to members of the Garcia lab, and to M. Akinyi and M. Algie for helpful discussions. We thank J. Kaur for assistance in the initial screen. Strains were provided by the *Caenorhabditis* Genetics Center (CGC). This work and J.T. were supported by the Academy of Finland Project Funding (309173), University of Helsinki Internal Funding and the Otto A. Malm Foundation. O.E. was supported by Finska Läkaresällskapet, J.T. and A.S.B by AICEP, Portuguese Trade & Investment Agency, Project Inov Contacto Program - international internships, 2017, 2018, co-financed by Portuguese and European funds.

## Author contributions

J.T. and S.M.D.A.G. designed the study. J.T. performed the experiments and analyzed the data. A.M.H. performed most of the qRT-PCR assays and together with A.B. assisted with the motility assays. J.T collected the samples and A.O. performed the lipid extraction and data analysis. O.E. and J.T performed the mitochondrial respiration assays, and O.E. assisted on the data analysis. B.B. set up the cell system with J.T., B.B. also contributed to cell data analysis, and provided key insight into RQC components. S.M.D.A.G. wrote the manuscript. B.B., J.T. and S.M.D.A.G. edited the manuscript.

## Abbreviations

ctl: catalase
CoQ: Coenzyme Q
DM1: myotonic dystrophy type 1
DMPK: dystrophia myotonica-protein kinase gene
Egl phenotype: egg-laying defective phenotype
EtOH: ethanol
ETC: electron transport chain
EV: empty vector (control RNAi)
GD: *E. coli* [*ubiG*::Kan, *zei*::Tn*10*dTet]
HT115: *E. coli* [F-, mcrA, mcrB, IN(rrnD-rrnE)1, rnc14::Tn10(DE3 lysogen: lacUV5 promoter -T7 polymerase]
LC-MS: liquid chromatography coupled to mass spectrometry
mtDNA: mitochondrial DNA
MBL-1: muscleblind splicing regulator homolog
MBNL: muscleblind-like splicing regulator
NAC: N-acetylcysteine
OCR: oxygen consumption rate
OP50: *E. coli* strain
RNAi: RNA interference
OXPHOS: oxidative phosphorylation
ROS: reactive oxygen species
RQC: ribosome-associated quality control
SOD: superoxide dismutase
SRC: spare respiratory capacity
UPRmt: mitochondrial unfolded protein response

## References

Adereth, Y., V. Dammai, N. Kose, R. Li, and T. Hsu. 2005. RNA-dependent integrin alpha3 protein localization regulated by the Muscleblind-like protein MLP1. Nat Cell Biol. 7:1240–1247.

Andrew, S.E., Y.P. Goldberg, B. Kremer, H. Telenius, J. Theilmann, S. Adam, E. Starr, F. Squitieri, B. Lin, M.A. Kalchman, and et al. 1993. The relationship between trinucleotide (CAG) repeat length and clinical features of Huntington’s disease. Nat Genet. 4:398–403.

Batra, R., K. Charizanis, M. Manchanda, A. Mohan, M. Li, D.J. Finn, M. Goodwin, C. Zhang, K. Sobczak, C.A. Thornton, and M.S. Swanson. 2014. Loss of MBNL leads to disruption of developmentally regulated alternative polyadenylation in RNA-mediated disease. Mol Cell. 56:311–322.

Batra, R., M. Manchanda, and M.S. Swanson. 2015. Global insights into alternative polyadenylation regulation. RNA Biol. 12:597–602.

Bratic, I., J. Hench, and A. Trifunovic. 2010. Caenorhabditis elegans as a model system for mtDNA replication defects. Methods. 51:437–443.

Brenner, S. 1974. The genetics of Caenorhabditis elegans. Genetics. 77:71–94.

Brook, J.D., M.E. McCurrach, H.G. Harley, A.J. Buckler, D. Church, H. Aburatani, K. Hunter, V.P. Stanton, J.P. Thirion, T. Hudson, and et al. 1992. Molecular basis of myotonic dystrophy: expansion of a trinucleotide (CTG) repeat at the 3’ end of a transcript encoding a protein kinase family member. Cell. 69:385.

Brown, R.E., and C.H. Freudenreich. 2021. Structure-forming repeats and their impact on genome stability. Curr Opin Genet Dev. 67:41–51.

Cajka, T., and O. Fiehn. 2016. Increasing lipidomic coverage by selecting optimal mobile-phase modifiers in LC–MS of blood plasma. 12:34.

Chau, A., and A. Kalsotra. 2015. Developmental insights into the pathology of and therapeutic strategies for DM1: Back to the basics. Dev Dyn. 244:377–390.

Chen, H., M. Vermulst, Y.E. Wang, A. Chomyn, T.A. Prolla, J.M. McCaffery, and D.C. Chan. 2010. Mitochondrial fusion is required for mtDNA stability in skeletal muscle and tolerance of mtDNA mutations. Cell. 141:280–289.

Conti, B. 2008. Considerations on temperature, longevity and aging. Cell Mol Life Sci. 65:1626–1630.

Cornaglia, M., L. Mouchiroud, A. Marette, S. Narasimhan, T. Lehnert, V. Jovaisaite, J. Auwerx, and M.A. Gijs. 2015. An automated microfluidic platform for C. elegans embryo arraying, phenotyping, and long-term live imaging. Sci Rep. 5:10192.

Crane, F.L., and P. Navas. 1997. The diversity of coenzyme Q function. Mol Aspects Med. 18 Suppl:S1–6.

Du, H., M.S. Cline, R.J. Osborne, D.L. Tuttle, T.A. Clark, J.P. Donohue, M.P. Hall, L. Shiue, M.S. Swanson, C.A. Thornton, and M. Ares, Jr. 2010. Aberrant alternative splicing and extracellular matrix gene expression in mouse models of myotonic dystrophy. Nat Struct Mol Biol. 17:187–193.

Dutton, P.L., Ohnishi, T., Darrouzet, E., Leonard, M.A., Sharp, R. F., Gibney, B.., Daldal, F., and Moser, C. C. 2000. Coenzyme Q: Molecular Mechanims in Health and Disease. CRC Press, Boca Raton, FL.

Fernandez-Ayala, D.J., G. Lopez-Lluch, M. Garcia-Valdes, A. Arroyo, and P. Navas. 2005. Specificity of coenzyme Q10 for a balanced function of respiratory chain and endogenous ubiquinone biosynthesis in human cells. Biochim Biophys Acta. 1706:174–183.

Garcia, S.M., Y. Tabach, G.F. Lourenco, M. Armakola, and G. Ruvkun. 2014. Identification of genes in toxicity pathways of trinucleotide-repeat RNA in C. elegans. Nat Struct Mol Biol. 21:712–720.

Goodwin, M., A. Mohan, R. Batra, K.Y. Lee, K. Charizanis, F.J. Fernandez Gomez, S. Eddarkaoui, N. Sergeant, L. Buee, T. Kimura, H.B. Clark, J. Dalton, K. Takamura, S.M. Weyn-Vanhentenryck, C. Zhang, T. Reid, L.P. Ranum, J.W. Day, and M.S. Swanson. 2015. MBNL Sequestration by Toxic RNAs and RNA Misprocessing in the Myotonic Dystrophy Brain. Cell Rep. 12:1159–1168.

Gramegna, L.L., M.P. Giannoccaro, D.N. Manners, C. Testa, S. Zanigni, S. Evangelisti, C. Bianchini, F. Oppi, R. Poda, P. Avoni, R. Lodi, R. Liguori, and C. Tonon. 2018. Mitochondrial dysfunction in myotonic dystrophy type 1. Neuromuscul Disord. 28:144–149.

Harper, P. 2001. Myotonic dystrophy. Muscle & Nerve. 25:926.

Heo, J.M., N. Livnat-Levanon, E.B. Taylor, K.T. Jones, N. Dephoure, J. Ring, J. Xie, J.L. Brodsky, F. Madeo, S.P. Gygi, K. Ashrafi, M.H. Glickman, and J. Rutter. 2010. A stress-responsive system for mitochondrial protein degradation. Mol Cell. 40:465–480.

Ho, T.H., B.N. Charlet, M.G. Poulos, G. Singh, M.S. Swanson, and T.A. Cooper. 2004. Muscleblind proteins regulate alternative splicing. EMBO J. 23:3103–3112.

Izawa, T., S.H. Park, L. Zhao, F.U. Hartl, and W. Neupert. 2017. Cytosolic Protein Vms1 Links Ribosome Quality Control to Mitochondrial and Cellular Homeostasis. Cell. 171:890–903 e818.

Joazeiro, C.A.P. 2019. Mechanisms and functions of ribosome-associated protein quality control. Nat Rev Mol Cell Biol. 20:368–383.

Jonassen, T., B.N. Marbois, K.F. Faull, C.F. Clarke, and P.L. Larsen. 2002. Development and fertility in Caenorhabditis elegans clk-1 mutants depend upon transport of dietary coenzyme Q8 to mitochondria. J Biol Chem. 277:45020–45027.

Kamath, R.S., M. Martinez-Campos, P. Zipperlen, A.G. Fraser, and J. Ahringer. 2001. Effectiveness of specific RNA-mediated interference through ingested double-stranded RNA in Caenorhabditis elegans. Genome Biol. 2:RESEARCH0002.

Kanazawa, T., M.D. Zappaterra, A. Hasegawa, A.P. Wright, E.D. Newman-Smith, K.F. Buttle, K. McDonald, C.A. Mannella, and A.M. van der Bliek. 2008. The C. elegans Opa1 homologue EAT-3 is essential for resistance to free radicals. PLoS Genet. 4:e1000022.

Koopman, M., H. Michels, B.M. Dancy, R. Kamble, L. Mouchiroud, J. Auwerx, E.A. Nollen, and R.H. Houtkooper. 2016. A screening-based platform for the assessment of cellular respiration in Caenorhabditis elegans. Nat Protoc. 11:1798–1816.

Lander, E.S., L.M. Linton, B. Birren, C. Nusbaum, M.C. Zody, J. Baldwin, K. Devon, K. Dewar, M. Doyle, W. FitzHugh, R. Funke, D. Gage, K. Harris, A. Heaford, J. Howland, L. Kann, J. Lehoczky, R. LeVine, P. McEwan, K. McKernan, J. Meldrim, J.P. Mesirov, C. Miranda, W. Morris, J. Naylor, C. Raymond, M. Rosetti, R. Santos, A. Sheridan, C. Sougnez, Y. Stange-Thomann, N. Stojanovic, A. Subramanian, D. Wyman, J. Rogers, J. Sulston, R. Ainscough, S. Beck, D. Bentley, J. Burton, C. Clee, N. Carter, A. Coulson, R. Deadman, P. Deloukas, A. Dunham, I. Dunham, R. Durbin, L. French, D. Grafham, S. Gregory, T. Hubbard, S. Humphray, A. Hunt, M. Jones, C. Lloyd, A. McMurray, L. Matthews, S. Mercer, S. Milne, J.C. Mullikin, A. Mungall, R. Plumb, M. Ross, R. Shownkeen, S. Sims, R.H. Waterston, R.K. Wilson, L.W. Hillier, J.D. McPherson, M.A. Marra, E.R. Mardis, L.A. Fulton, A.T. Chinwalla, K.H. Pepin, W.R. Gish, S.L. Chissoe, M.C. Wendl, K.D. Delehaunty, T.L. Miner, A. Delehaunty, J.B. Kramer, L.L. Cook, R.S. Fulton, D.L. Johnson, P.J. Minx, S.W. Clifton, T. Hawkins, E. Branscomb, P. Predki, P. Richardson, S. Wenning, T. Slezak, N. Doggett, J.F. Cheng, A. Olsen, S. Lucas, C. Elkin, E. Uberbacher, M. Frazier, et al. 2001. Initial sequencing and analysis of the human genome. Nature. 409:860–921.

Liesa, M., and O.S. Shirihai. 2013. Mitochondrial dynamics in the regulation of nutrient utilization and energy expenditure. Cell Metab. 17:491–506.

Mahadevan, M., C. Tsilfidis, L. Sabourin, G. Shutler, C. Amemiya, G. Jansen, C. Neville, M. Narang, J. Barcelo, K. O’Hoy, and et al. 1992. Myotonic dystrophy mutation: an unstable CTG repeat in the 3’ untranslated region of the gene. Science. 255:1253–1255.

Mankodi, A., E. Logigian, L. Callahan, C. McClain, R. White, D. Henderson, M. Krym, and C.A. Thornton. 2000. Myotonic dystrophy in transgenic mice expressing an expanded CUG repeat. Science. 289:1769–1773.

Masuda, A., H.S. Andersen, T.K. Doktor, T. Okamoto, M. Ito, B.S. Andresen, and K. Ohno. 2012. CUGBP1 and MBNL1 preferentially bind to 3’ UTRs and facilitate mRNA decay. Sci Rep. 2:209.

Miller, J.W., C.R. Urbinati, P. Teng-Umnuay, M.G. Stenberg, B.J. Byrne, C.A. Thornton, and M.S. Swanson. 2000. Recruitment of human muscleblind proteins to (CUG)(n) expansions associated with myotonic dystrophy. EMBO J. 19:4439–4448.

Mitchell, P. 1975. Protonmotive redox mechanism of the cytochrome b-c1 complex in the respiratory chain: protonmotive ubiquinone cycle. FEBS Lett. 56:1–6.

Nemoto, Y., and P. De Camilli. 1999. Recruitment of an alternatively spliced form of synaptojanin 2 to mitochondria by the interaction with the PDZ domain of a mitochondrial outer membrane protein. EMBO J. 18:2991–3006.

Norris, A.D., X. Gracida, and J.A. Calarco. 2017. CRISPR-mediated genetic interaction profiling identifies RNA binding proteins controlling metazoan fitness. Elife. 6.

Ono, S., H. Kurisaki, K. Inouye, and T. Mannen. 1986. “Ragged-red” fibres in myotonic dystrophy. J Neurol Sci. 74:247–255.

Paulson, H. 2018. Repeat expansion diseases. Handb Clin Neurol. 147:105–123.

Pellegrino, R.M., A. Di Veroli, A. Valeri, L. Goracci, and G. Cruciani. 2014. LC/MS lipid profiling from human serum: a new method for global lipid extraction. Anal Bioanal Chem. 406:7937–7948.

Rea, S.L., N. Ventura, and T.E. Johnson. 2007. Relationship between mitochondrial electron transport chain dysfunction, development, and life extension in Caenorhabditis elegans. PLoS Biol. 5:e259.

Rodriguez-Aguilera, J.C., C. Asencio, M. Ruiz-Ferrer, J. Vela, and P. Navas. 2003. Caenorhabditis elegans ubiquinone biosynthesis genes. Biofactors. 18:237–244.

Rudnicki, D.D., S.E. Holmes, M.W. Lin, C.A. Thornton, C.A. Ross, and R.L. Margolis. 2007. Huntington’s disease--like 2 is associated with CUG repeat-containing RNA foci. Ann Neurol. 61:272–282.

Sasagawa, N., E. Ohno, Y. Kino, Y. Watanabe, and S. Ishiura. 2009. Identification of Caenorhabditis elegans K02H8.1 (CeMBL), a functional ortholog of mammalian MBNL proteins. J Neurosci Res. 87:1090–1097.

Swinnen, B., W. Robberecht, and L. Van Den Bosch. 2020. RNA toxicity in non-coding repeat expansion disorders. EMBO J. 39:e101112.

Takahashi, M., M. Ogawara, T. Shimizu, and T. Shirasawa. 2012. Restoration of the behavioral rates and lifespan in clk-1 mutant nematodes in response to exogenous coenzyme Q(10). Exp Gerontol. 47:276–279.

Thomas, J.D., L.J. Sznajder, O. Bardhi, F.N. Aslam, Z.P. Anastasiadis, M.M. Scotti, I. Nishino, M. Nakamori, E.T. Wang, and M.S. Swanson. 2017. Disrupted prenatal RNA processing and myogenesis in congenital myotonic dystrophy. Genes Dev. 31:1122–1133.

Timmons, L., D.L. Court, and A. Fire. 2001. Ingestion of bacterially expressed dsRNAs can produce specific and potent genetic interference in Caenorhabditis elegans. Gene. 263:103–112.

Tokgozoglu, L.S., T. Ashizawa, A. Pacifico, R.M. Armstrong, H.F. Epstein, and W.A. Zoghbi. 1995. Cardiac involvement in a large kindred with myotonic dystrophy. Quantitative assessment and relation to size of CTG repeat expansion. JAMA. 274:813–819.

Tran, U.C., and C.F. Clarke. 2007. Endogenous synthesis of coenzyme Q in eukaryotes. Mitochondrion. 7 Suppl:S62-71.

Tsang, W.Y., and B.D. Lemire. 2002. Mitochondrial genome content is regulated during nematode development. Biochem Biophys Res Commun. 291:8–16.

Turner, C., and D. Hilton-Jones. 2010. The myotonic dystrophies: diagnosis and management. J Neurol Neurosurg Psychiatry. 81:358–367.

van der Bliek, A.M., Q. Shen, and S. Kawajiri. 2013. Mechanisms of mitochondrial fission and fusion. Cold Spring Harb Perspect Biol. 5.

Vita, G., A. Toscano, A. Prelle, A. Bordoni, N. Checcarelli, N. Bresolin, A. Baradello, and C. Messina. 1993. Muscle mitochondria investigation in myotonic dystrophy. Eur Neurol. 33:423–427.

Wagner, S.D., A.J. Struck, R. Gupta, D.R. Farnsworth, A.E. Mahady, K. Eichinger, C.A. Thornton, E.T. Wang, and J.A. Berglund. 2016. Dose-Dependent Regulation of Alternative Splicing by MBNL Proteins Reveals Biomarkers for Myotonic Dystrophy. PLoS Genet. 12:e1006316.

Wang, E.T., N.A. Cody, S. Jog, M. Biancolella, T.T. Wang, D.J. Treacy, S. Luo, G.P. Schroth, D.E. Housman, S. Reddy, E. Lecuyer, and C.B. Burge. 2012. Transcriptome-wide regulation of pre-mRNA splicing and mRNA localization by muscleblind proteins. Cell. 150:710–724.

Youle, R.J., and A.M. van der Bliek. 2012. Mitochondrial fission, fusion, and stress. Science. 337:1062–1065.

Yu, R., U. Lendahl, M. Nister, and J. Zhao. 2020. Regulation of Mammalian Mitochondrial Dynamics: Opportunities and Challenges. Front Endocrinol (Lausanne). 11:374.

