## Supplementary Figures for "RNA nucleotide repeats induce mitochondrial dysfunction and the ribosome-associated quality control"

### Supplementary figure 1

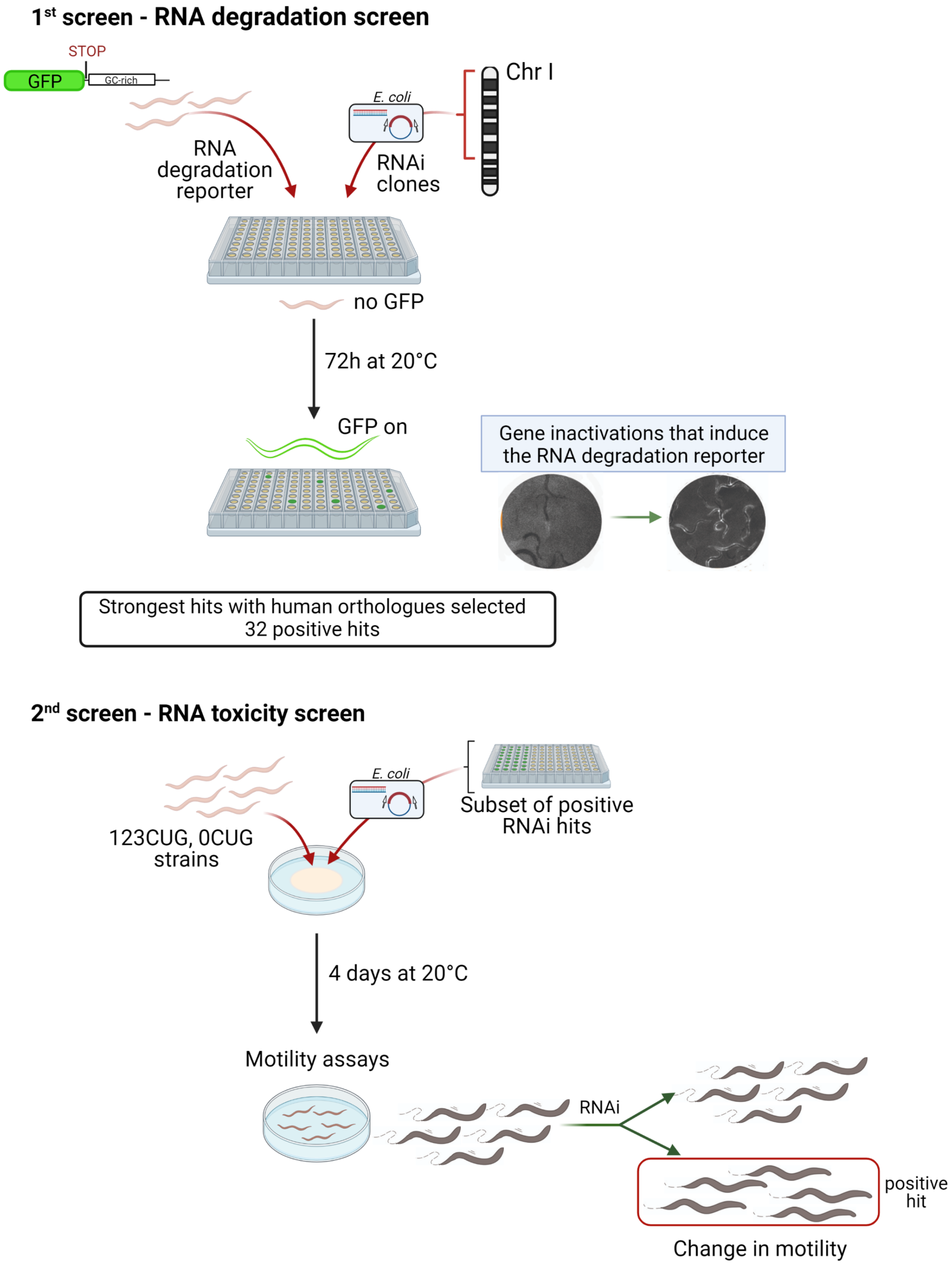

### Supplementary figure 2

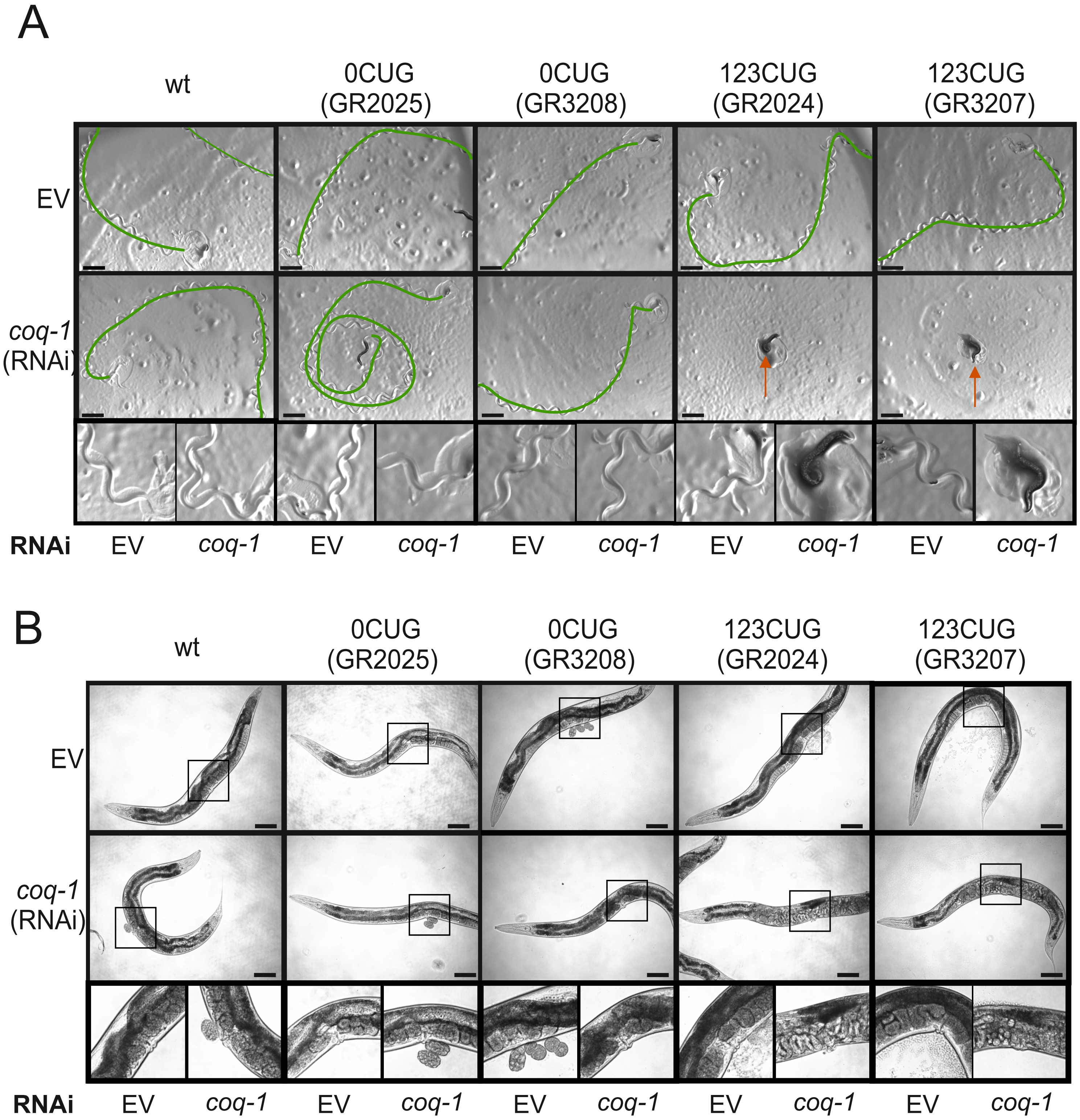

### Supplementary figure 3

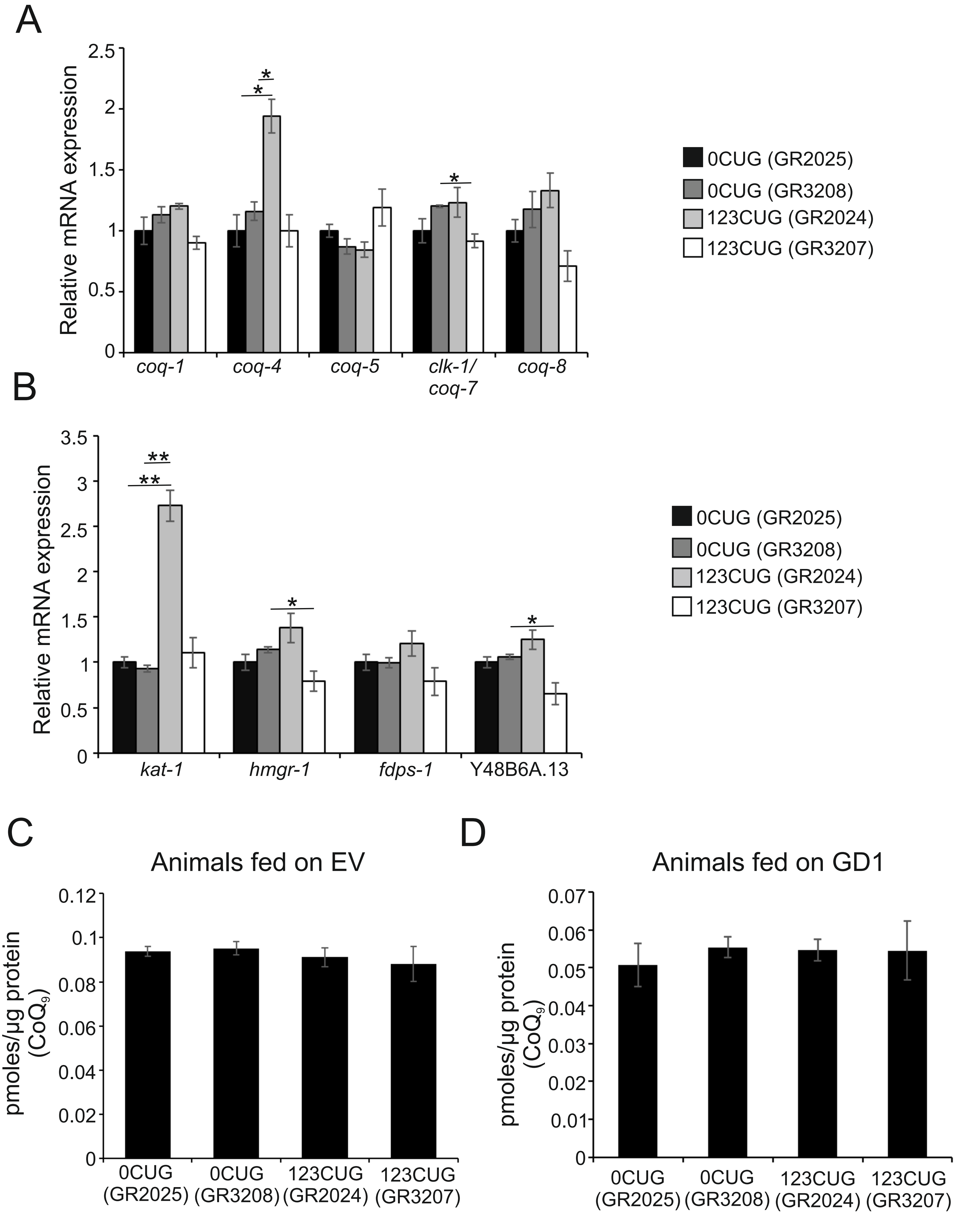

### Supplementary figure 4

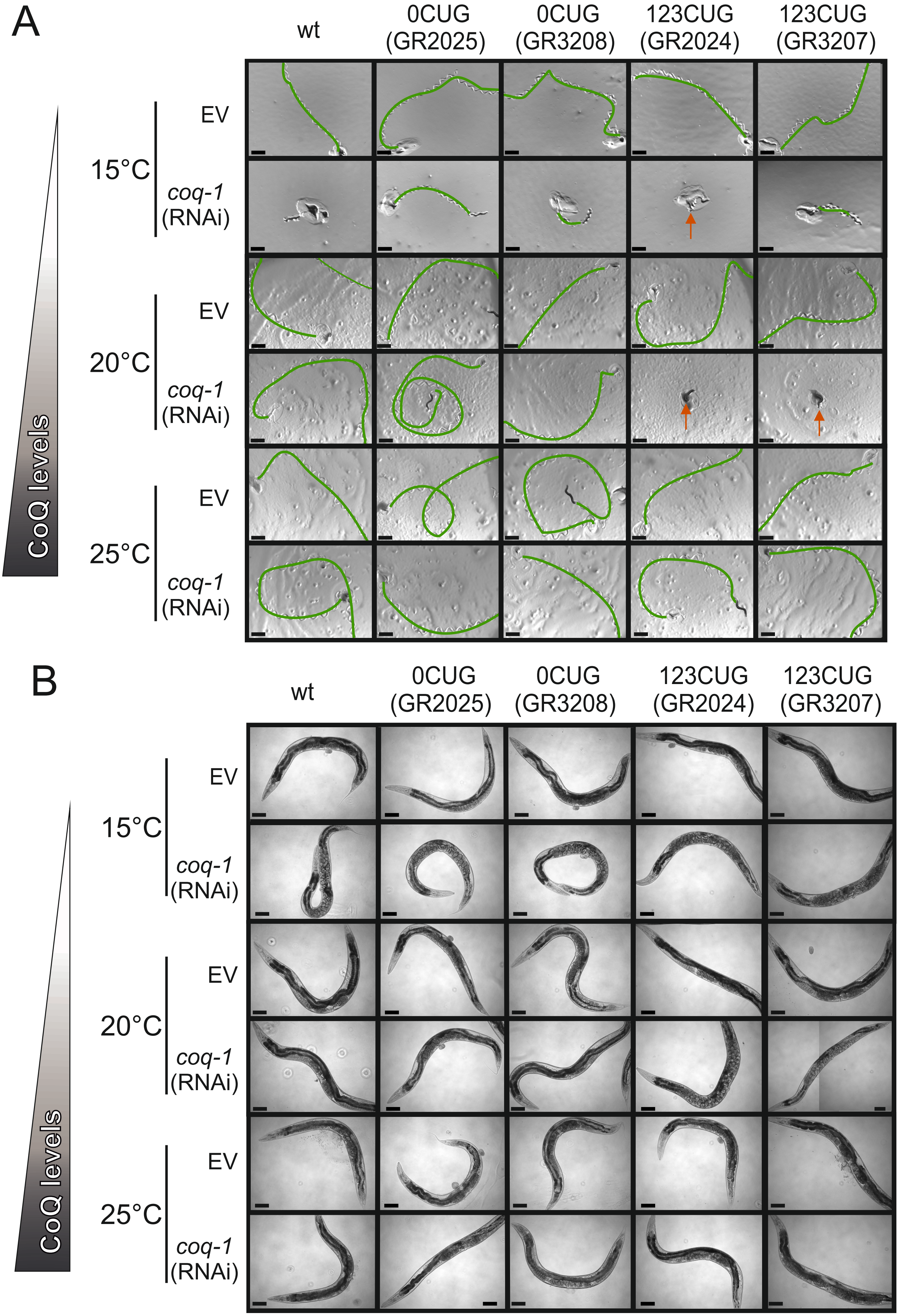

### Supplementary figure 5

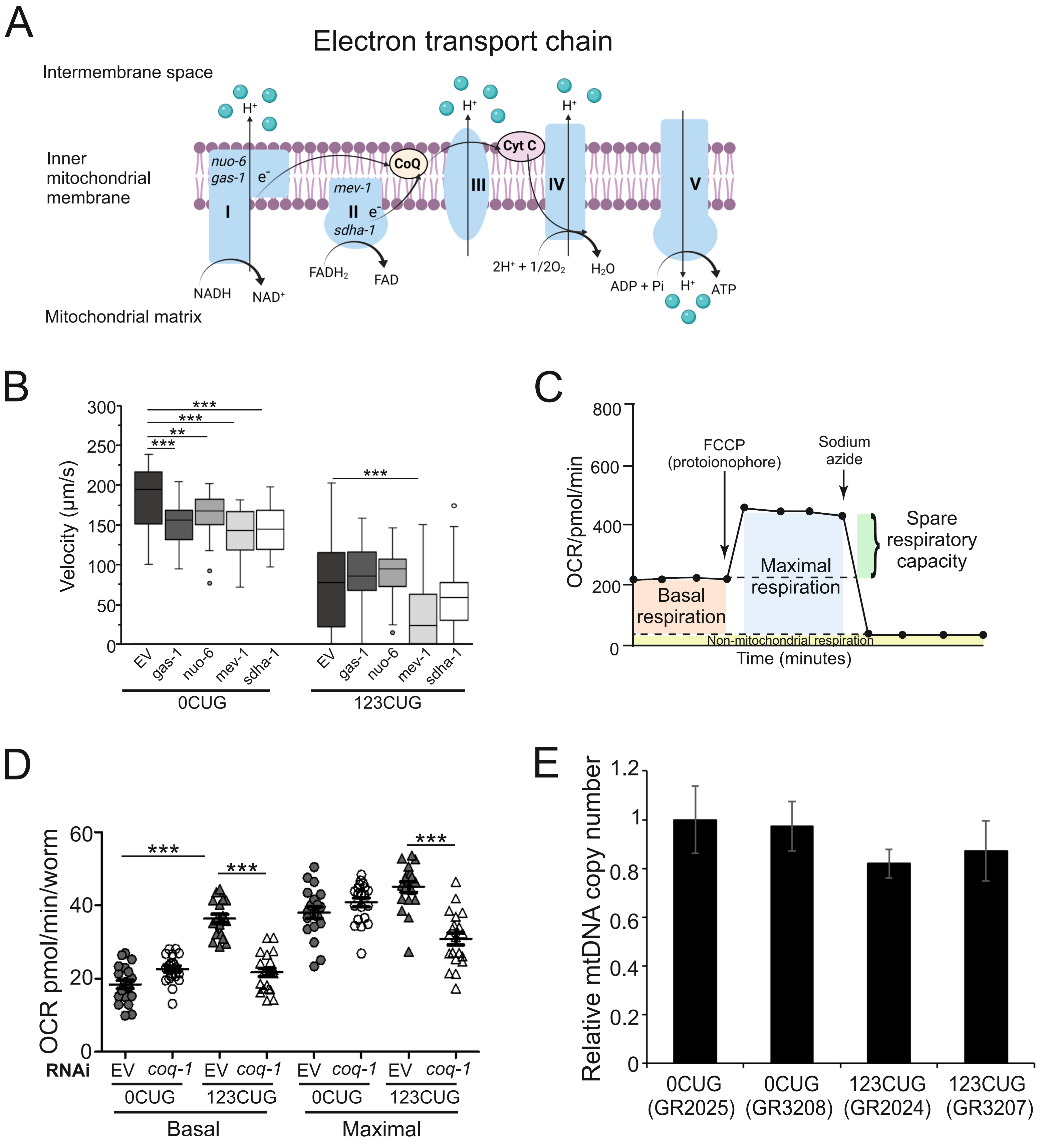

### Supplementary figure 6

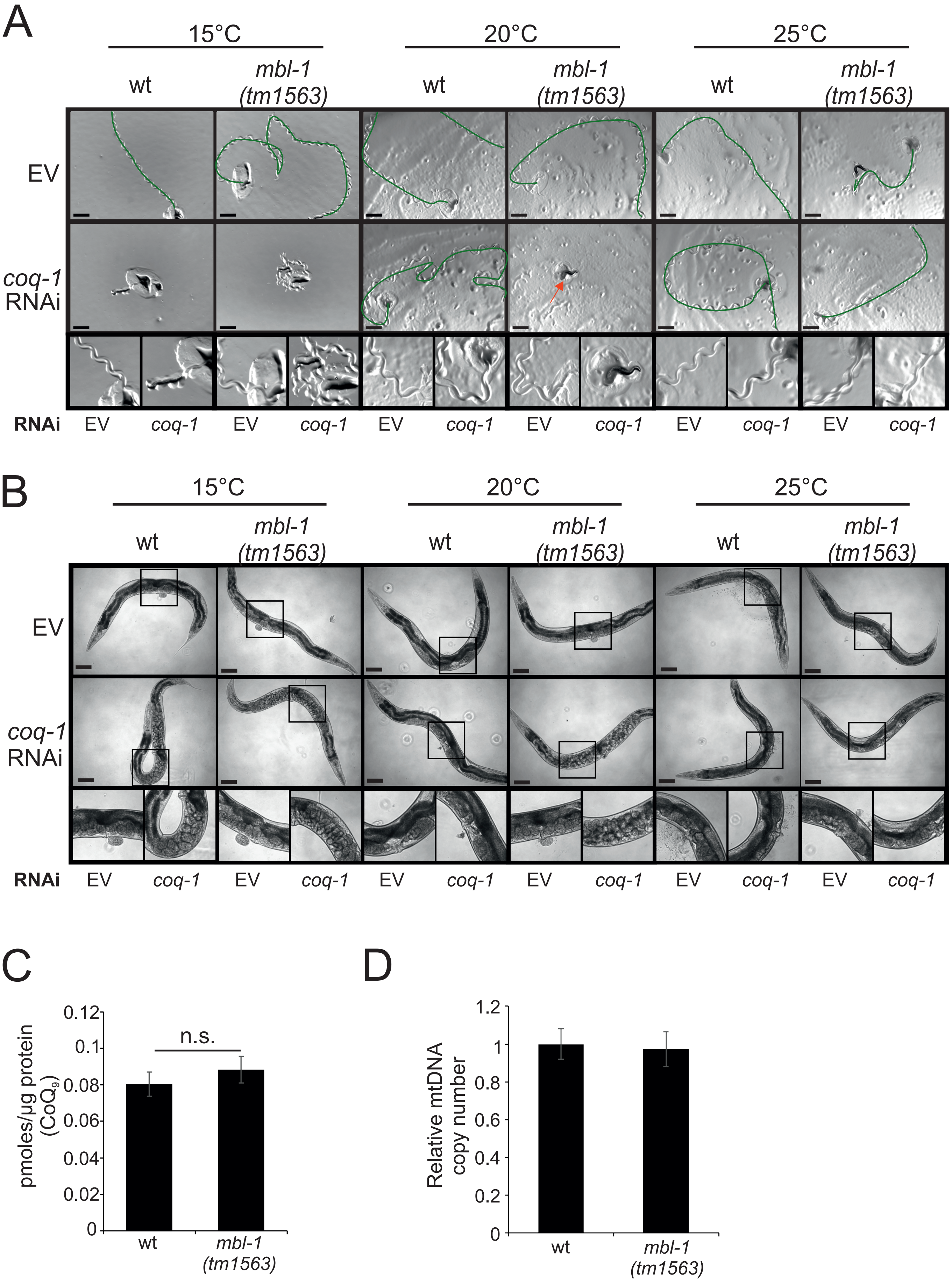

**Supplementary Figures Legends**

Supplementary Figure 1: Diagram of RNAi screens for the identification of modifiers of RNA repeat toxicity.

Two sequential RNAi screens were performed to identify modifiers of RNA repeat toxicity. 1st screen: a *C. elegans* RNA reporter strain (GR2173), expressing a GFP with a GC-rich 3'UTR, was used to analyze by RNAi two-thirds of Chr I and identify genes that function as potential modifiers of RNA degradation pathways. Genes were selected as positives that, when inactivated, interfered with the degradation of the reporter transcript, leading GR2173 animals to express a GFP fluorescent signal. 2nd screen: the strongest hits with human orthologues (32 clones) identified from the first screen were selected. These RNAi clones were used in motility assays with 123CUG and 0CUG strains to identify modifiers of CUG expanded toxicity. Genes were identified as positives for their ability, when inactivated, to modulate the motility defect of 123CUG animals without similarly affecting the 0CUG strain. For the detailed motility experimental procedure, see Methods. Image created with BioRender.com.

Supplementary Figure 2: Phenotypes observed upon inactivation of *coq-1* are repeat dependent.

(A – B) Representative DIC images of 0CUG (GR2025 and GR3208) and 123CUG (GR2024 and GR3207) strains fed EV and *coq-1* RNAis. (A) *coq-1* inactivation causes paralysis of 123CUG strains but does not affect the controls. Green line indicates visible tracks left by the animals, and arrow indicates the paralyzed animal. Scale bar, 1mm. (B) Presence of Egl phenotype in 123CUG when *coq-1* is inactivated. Scale bar, 100µm.

Supplementary Figure 3: Expression of CoQ biosynthetic factors is not affected in 123CUG animals. Analysis of mRNA relative expression of genes from (A) CoQ and (B) mevalonate pathways. $CoQ_9$ lipid levels of 0CUG (GR2025 and GR3208) and 123CUG (GR2024 and GR3207) strains fed (C) EV or (D) GD1 bacteria. Bars represent mean ± S.E.M. *P* value determined by two-tailed Student's *t*-test, *p<0.05, **p<0.005, ***p<0.0001.

Supplementary Figure 4: 123CUG animals show temperature-dependent rescue of RNA repeat toxicity

Representative DIC images of wild type (wt), 0CUG, and 123CUG strains fed EV and *coq-1* RNAis, at 15°C, 20°C, and 25°C. (A) *coq-1* inactivation causes paralysis in a temperature-dependent manner. 123CUG animals are more susceptible to *coq-1* inactivation than control strains.  Green line indicates visible tracks left by the animals and arrow indicates the paralyzed animals. Scale bar, 1mm. (B) Presence of Egl phenotype in 123CUG (GR2024 and GR3207) versus 0CUG (GR2025 and GR3208) and wt strains when *coq-1* is inactivated at 15°C, 20°C and 25°C. Scale bar, 100μm.

Supplementary Figure 5: 123CUG have dysfunctional mitochondria.

(A) Schematic representation of the electron transport chain (created with BioRender.com). During mitochondrial respiration, the electrons resulting from the conversion of NADH to $NAD^+$ (complex I) and $FADH_2$ to FAD (complex II) are transferred by CoQ to complex III. At the same time, complexes I, III and IV pump protons present in the mitochondrial matrix to the intermembrane space. These mechanisms produce a proton gradient that is used to generate ATP through complex V. (B) Velocity of 0CUG and 123CUG strains fed RNAi of mitochondrial components *gas-1*, *nuo-6* (complex I), *mev-1* and *sdha-1* (complex II). The RNAi was diluted with EV to the final concentration of 20% (*gas-1* and *nuo-6*) and 40% (*mev-1*).  Statistical significance was determined by Wilcoxon statistical test with Bonferroni correction. $^{**}p<0.005$; $^{***}p<0.0001$. n≥40 animals/condition. Box: $25^{th}$ to $75^{th}$ percentile; whiskers: 1.5 * interquartile range; line in the box: median; dots: outliers. (C) Representative oxygen consumption rates (OCR) profile of mitochondrial function assay. The mitochondrial uncoupler FCCP and the mitochondrial inhibitor sodium azide are used to determine maximal respiration and oxygen consumption from non-mitochondrial sites, respectively. (D) Effect of *coq-1* downregulation in the respiration of 0CUG and 123CUG animals. Two independent experiments were performed, n=10 wells/experiment. *P* value determined by two-way Anova coupled with

Bonferroni correction analysis; ***p<0.0001. (E) Relative mtDNA expression of 0CUG and 123CUG independent strains fed on empty vector RNAi. Four experiments were performed with at least three independent biological samples. Bars represent mean ± S.E.M. Statistical significance was determined by two-tailed Student's *t*-test.

Supplementary Figure 6: Mitochondrial dysregulation is dependent on MBL-1.

(A-B) Representative DIC images of wt and *mbl-1(tm1563)* animals fed EV and *coq-1* RNAi at 15°C, 20°C and 25°C. (A) *coq-1* inactivation causes paralysis in a temperature-dependent manner in *mbl-1* mutants. Green line indicates visible tracks left by the animals and arrow indicates the paralyzed animal. Scale bar, 1mm. (B) Presence of Egl phenotype in *mbl-1* mutants at 15°C and 20°C and in wt at 15°C. Scale bar, 100µm. (C) Analysis of $CoQ_9$ levels of wt and *mbl-1(tm1563)* animals fed on empty vector. (D) Relative mtDNA expression of wt and *mbl-1* mutants fed on empty vector RNAi. Two experiments were performed with at least three independent biological samples. (C-D) Bars represent mean ± S.E.M. Statistical significance was determined by two-tailed Student's *t*-test.
