## Supplementary Tables for "RNA nucleotide repeats induce mitochondrial dysfunction and the ribosome-associated quality control"

Supplementary table 1 – List of strains

| Strain | Genotype |
|---|---|
| Bristol N2 | wild type |
| FX01563 | *mbl-1(tm1563)* |
| GR2024 | mgIs64[*myo-3*p::gfp::3'utr123(CUG)] |
| GR2025 | mgIs65[*myo-3*p::gfp::3'utr0(CUG)] |
| GR2173 | mgIs71[p*myo-3*::gfp::3'utr(70%Gcinsert);p*myo-2*::(nls)mcherry] |
| GR3207 | mgIs84[*myo-3*p::gfp::3'utr123(CUG)] |
| GR3208 | mgIs85[*myo-3*p::gfp::3'utr0(CUG)] |
| GAR118 | iceEx42[*unc-54*p::mcherry::OMP25] |
| GAR126 | iceEx42[*unc-54*p::mcherry::OMP25]; mgIs64[*myo-3*p::gfp::3'utr123(CUG)] |
| GAR131 | iceEx42[*unc-54*p::mcherry::OMP25]; mgIs65[*myo-3*p::gfp::3'utr0(CUG)] |
| GAR138 | iceEx42[*unc-54*p::mcherry::OMP25]; *mbl-1(tm1563)]* |

Supplementary table 2 – List of primers

Sequence in bold and in bold and italic correspond to the sequences of mCherry and pPD49.26, respectively.

| qPCR | | |
|---|---|---|
| **Gene** | **Forward (5'→3')** | **Reverse (5'→3')** |
| *act-1* | TGCGACATTGATATCCGTAAGG | GGTGGTTCTCCGGAAAGAA |
| *nduo-1* | AGCGTCATTTATTGGGAAGAAGAC | AAGCTTGTGCTAATCCCATAAATGT |
| *cdc-42* | CTGCTGGACAGGAAGATTACG | CTCGGACATTCTCGAATGAAG |
| *pmp-3* | GTTCCCGTGTTCATCACTCAT | ACACCGTCGAGAAGCTGTAGA |
| *coq-1* | GATCCATACAGCGTCTCTAGTG | CGAGCAAGAATGAAATCTCCG |
| *coq-4* | TGGAAAGCGACTTCTTCTCGA | TGGAGATGTGTTCAAACGATCG |
| *coq-5* | ATGAGGCAGAGAAGGAGCAG | TGCAGGCCACCAACATAATA |
| *clk-1* | TGAAAGAACTCCTTGCCGAC | CCATTTGAGAGCCGAGTAGG |
| *coq-8* | TTTGCTAATCAGGATGTGACAATG | GTAGCTTTGAGTTTTGCGGC |
| *sod-1* | ACGCTTTACGGTCCAAACACT | CTTGGACTCTTCTGCCTTGTCT |
| *sod-2* | GTCACGAGATTATGCAACTTCATC | TCCATTGAACTTGAGAGCTGG |
| *sod-3* | TCAAAGGAGCTGATGGACAC | CAATATCCCAACCATCCCCAG |
| *ctl-1* | AGATGGTGCTGAACAGGAATG | GTAATGCGTGTCCGTGTAGG |
| *ctl-2* | CAAGATGGTGCTGAACAGAAATC | TGGAAATGAGTGTCGGTGTAC |
| *trx-2* | GTTGATTTCCACGCAGAATGG | TCCATTGCCAGTTCACCAG |
| *kat-1* | CATGGGAAACTGCGGAGAG | GATTTGACGTTCACTGGCAC |
| *hmgr-1* | GTCTTGCCGAAGTTATTGCG | TGAGTTGCTTTTCCAGAGTCG |
| *fdps-1* | GAGAAAGTGGAGAAAATTAAGCGG | GTCAGGAATCTCGGATATGCTC |
| Y48B6A.13 | GGCAGTTAATCGACACAGTTG | TGGATAGTTTGGAGCACAGC |
| **Cloning** | | |
| pPD49.26 | GATATCTGAGCTCCGCATCGGC | CTTGTACAGCTCGTCCATG |
| OMP25 | **GCATGGACGAGCTGTACAAGG**GCGACGGAGAGCCGAGT | ***CGATGCGGAGCTCAGATATC***GCTGGGTTAACTATTAGAGCTGCTTTC |

Supplementary table 3 – Percentages of mitochondrial fragmentation in *C. elegans*.

| **Strain, RNAi and treatment** | **Tubular (%)** | **Fragmented (%)** | **Very fragmented (%)** |
|---|---|---|---|
| **Figure 2E** | | | |
| wt | 78,2 | 13,4 | 8,4 |
| 0CUG | 78,8 | 13,6 | 7,6 |
| 123CUG | 49,0 | 30,2 | 20,8 |
| **Figure 2F** | | | |
| 0CUG EV | 79,5 | 7,7 | 12,8 |
| 0CUG *coq-1* | 46,5 | 34,9 | 18,6 |
| 0CUG *coq-1* EtOH | 58,0 | 20,0 | 22,0 |
| 0CUG *coq-1* COQ | 56,4 | 18,2 | 25,5 |
| 123CUG EV | 46,3 | 24,1 | 29,6 |
| 123CUG *coq-1* | 22,0 | 36,6 | 41,5 |
| 123CUG *coq-1* EtOH | 42,6 | 23,5 | 33,8 |
| 123CUG *coq-1* COQ | 68,8 | 14,1 | 17,2 |
| **Figure 4E** | | | |
| wt | 73,1 | 16,2 | 10,7 |
| wt EtOH | 41,3 | 29,5 | 29,3 |
| wt CoQ | 32,8 | 37,7 | 29,5 |
| *mbl-1* | 36,1 | 42,5 | 21,4 |
| *mbl-1* EtOH | 34,7 | 36,6 | 28,6 |
| *mbl-1* CoQ | 63,8 | 25,5 | 10,7 |

Supplementary table 4 – Percentages of mitochondrial fragmentation in HeLa cells.

| **Strain, RNAi and treatment** | **Tubular (%)** | **Intermediate (%)** | **Fragmented (%)** | **Aggregated (%)** |
|---|---|---|---|---|
| **Figure 3C** | | | | |
| Non-transfected | 40,9 | 41,4 | 11,6 | 6,1 |
| GFP | 23,1 | 58,4 | 13,3 | 5,1 |
| 5CTG | 15,2 | 65,4 | 15,0 | 4,3 |
| 100CTG | 4,2 | 61,6 | 25,3 | 8,9 |
| 200CTG | 3,0 | 52,8 | 22,4 | 21,8 |

Supplementary table 5 – Statistical analysis

| **Strain, RNAi and treatment** | **p-value** | **Bonferroni correction** |
|---|---|---|
| **Figure 1A - Wilcoxon test** | | |
| 0CUG EV vs 0CUG *coq-1* | <0,0001 | |
| 0GUG EV vs 123CUG EV | <0,0001 | p < 0,025 |
| 123CUG EV vs 123CUG *coq-1* | <0,0001 | |
| **Figure 1C - *t*-test** | | |
| 0CUG EV vs 123CUG EV | 0,82 | n.a. |
| 0CUG *coq-1* vs 123CUG *coq-1* | 0,07 | |

| | | | |
|---|---|---|---|
| **Figure 1D - Wilcoxon test** | | | |
| 0CUG vs 0CUG EtOH | | 0,99 | p < 0,017 |
| 0CUG EtOH vs 0CUG CoQ | | 0,88 | |
| 0CUG EtOH vs 123CUG EtOH | | <0,0001 | |
| 123CUG vs 123CUG EtOH | | 0,35 | |
| 123CUG EtOH vs 123CUG CoQ | | 1,50E-03 | |
| 0CUG vs 123CUG | | <0,0001 | |
| **Figure 2A - *t*-test** | | | |
| Basal | 0CUG vs 123CUG | <0,0001 | p < 0,025 |
| Maximal | 0CUG vs 123CUG | <0,0001 | |
| **Figure 2B - *t*-test** | | | |
| 0CUG vs 123CUG | | 9,42E-05 | n.a. |
| **Figure 2C - 2-way Anova with bonferroni correction** | | | |
| Basal | 0CUG- vs 0CUG EtOH | 0,06 | n.a. |
| | 0CUG- vs 0CUG CoQ | 1,00 | |
| | 0CUG EtOH vs 0CUG CoQ | 0,72 | |
| | 0CUG- vs 123CUG- | 6,93E-17 | |
| | 0CUG EtOH vs 123CUG EtOH | 1,75E-16 | |
| | 0CUG- vs 123CUG CoQ | 8,80E-09 | |
| | 0CUG EtOH vs 123CUG CoQ | 5,00E-06 | |
| | 0CUG CoQ vs 123CoQ | 2,23E-10 | |
| | 123CUG- vs 123CUG EtOH | 1,00 | |
| | 123CUG- vs 123CUG CoQ | 5,40E-05 | |
| | 123CUG EtOH vs 123CUG CoQ | 1,30E-04 | |
| Maximal | 0CUG- vs 0CUG EtOH | 1,00 | |
| | 0CUG- vs 0CUG CoQ | 0,53 | |
| | 0CUG EtOH vs 0CUG CoQ | 0,72 | |
| | 0CUG- vs 123CUG- | 1,10E-05 | |
| | 0CUG EtOH vs 123CUG EtOH | 2,40E-05 | |
| | 0CUG- vs 123CUG CoQ | 0,54 | |
| | 0CUG EtOH vs 123CUG CoQ | 0,25 | |
| | 0CUG CoQ vs 123CoQ | 0,01 | |
| | 123CUG- vs 123CUG EtOH | 0,51 | |
| | 123CUG- vs 123CUG CoQ | 1,08E-04 | |
| | 123CUG EtOH vs 123CUG CoQ | 0,01 | |
| **Figure 2E - chi-square test** | | | |
| wt vs 0CUG | | 0,99 | p < 0,017 |
| 0CUG vs 123CUG | | 88,0E-6 | |
| **Figure 2F - chi-square test** | | | |
| 0CUG | EV vs *coq-1* | <0,00001 | p < 0,0125 |
| | *coq-1*vs *coq-1* EtOH | 0,07 | |
| | *coq-1* ctrl vs *coq-1* CoQ | 0,85 | |
| 123CUG | EV vs *coq-1* | 1,55E-03 | |
| | *coq-1*vs *coq-1* EtOH | 4,74E-03 | |
| | *coq-1* EtoH vs *coq-1* CoQ | 9,63E-04 | |
| 0CUG EV vs 123CUG EV | | <0,00001 | |
| **Figure 3C - chi-square test** | | | |
| Non transfected vs GFP | | 4,64E-02 | p < 0,01 |
| GFP vs 5CTG | | 0,20 | |

| | | | |
|---|---|---|---|
| 5CTG vs 100CTG | | 1,25E-02 | |
| 5CTG vs 200CTG | | 4,00E-05 | |
| 100CTG vs 200CTG | | 0,09 | |
| **Figure 3E - Wilcoxon** | | | |
| 0CUG | EV vs *fzo-1* | 0,12 | p < 0,0125 |
| | EV vs *drp-1* | 0,26 | |
| | EV vs *eat-3* | 0,06 | |
| 123CUG | EV vs *fzo-1* | 0,23 | |
| | EV vs *drp-1* | 0,30 | |
| | EV vs *eat-3* | <0,0001 | |
| 0CUG EV vs 123CUG EV | | <0,0001 | |
| **Figure 4A - Wilcoxon** | | | |
| wt vs *mbl-1* | | <0,0001 | p < 0,017 |
| wt vs wt EtOH | | <0,0001 | |
| wt EtOH vs wt CoQ | | <0,0001 | |
| *mbl-1 vs mbl-1* EtOH | | <0,0001 | |
| wt EtOH vs *mbl-1* EtOH | | <0,0001 | |
| *mbl-1* EtOH *vs mbl-1 CoQ* | | 7,10E-03 | |
| **Figure 4B - Wilcoxon** | | | |
| wt | EV vs *gas-1* 20% | <0,0001 | p < 0,01 |
| | EV vs *nuo-6* 20% | <0,0001 | |
| | EV vs *mev-1* 40% | 0,11 | |
| | EV vs *sdha-1* | <0,0001 | |
| *mbl-1* | EV vs *gas-1* 20% | 0,55 | |
| | EV vs *nuo-6* 20% | 0,40 | |
| | EV vs *mev-1* 40% | 0,01 | |
| | EV vs *sdha-1* | 0,25 | |
| wt EV vs *mbl-1* EV | | <0,0001 | |
| **Figure 4C - *t*-test** | | | |
| Basal | wt vs *mbl-1* | < 0,0001 | p < 0,025 |
| FCCP | wt vs *mbl-1* | 0,10 | |
| **Figure 4D - *t*-test** | | | |
| 0CUG vs 123CUG | | 0,029 | n.a. |
| **Figure 4E - chi-square test** | | | |
| wt | EV vs EtOH | 3,0E-05 | p < 0,017 |
| | EtOH vs CoQ | 0,36 | |
| *mbl-1* | EV vs EtOH | 0,42 | |
| | EtOH vs CoQ | 9,50E-05 | |
| wt EV vs *mbl-1* EV | | <0,00001 | |
| wt EV vs *mbl-1* CoQ | | 0,23 | |
| **Figure 5A - Wilcoxon** | | | |
| 0CUG | EV vs *vms-1* | 0,9595 | p < 0,017 |
| | EV vs *cdc-48.2* | <0,0001 | |
| 123CUG | EV vs *vms-1* | <0,0001 | |
| | EV vs *cdc-48.2* | <0,0001 | |
| 0CUG EV vs 123CUG EV | | <0,0001 | |
| **Figure 5B - Wilcoxon** | | | |
| wt EV vs wt *cdc-48.2* | | 0,0011 | p < 0,025 |

| | | | |
|---|---|---|---|
| wt EV vs *mbl-1* EV | | <0,0001 | |
| *mbl-1* EV vs *mbl-1 cdc-48.2* | | <0,0001 | |
| **Supplementary Figure 3A - *t*-test** | | | |
| *coq-1* | 0CUG(GR2025) vs 0CUG(GR3208) | 0,38 | n.a. |
| | 0CUG(GR2025) vs 123CUG(GR2024) | 0,21 | |
| | 0CUG(GR2025) vs 123CUG(GR3207) | 0,49 | |
| | 0CUG(GR3208) vs 123CUG(GR2024) | 0,41 | |
| | 0CUG(GR3208) vs 123CUG(GR3207) | 0,06 | |
| *coq-4* | 0CUG(GR2025) vs 0CUG(GR3208) | 0,35 | |
| | 0CUG(GR2025) vs 123CUG(GR2024) | 0,008 | |
| | 0CUG(GR2025) vs 123CUG(GR3207) | 0,99 | |
| | 0CUG(GR3208) vs 123CUG(GR2024) | 0,014 | |
| | 0CUG(GR3208) vs 123CUG(GR3207) | 0,36 | |
| *coq-5* | 0CUG(GR2025) vs 0CUG(GR3208) | 0,19 | |
| | 0CUG(GR2025) vs 123CUG(GR2024) | 0,13 | |
| | 0CUG(GR2025) vs 123CUG(GR3207) | 0,34 | |
| | 0CUG(GR3208) vs 123CUG(GR2024) | 0,78 | |
| | 0CUG(GR3208) vs 123CUG(GR3207) | 0,16 | |
| *clk-1/coq-7* | 0CUG(GR2025) vs 0CUG(GR3208) | 0,17 | |
| | 0CUG(GR2025) vs 123CUG(GR2024) | 0,20 | |
| | 0CUG(GR2025) vs 123CUG(GR3207) | 0,51 | |
| | 0CUG(GR3208) vs 123CUG(GR2024) | 0,81 | |
| | 0CUG(GR3208) vs 123CUG(GR3207) | 0,03 | |
| *coq-8* | 0CUG(GR2025) vs 0CUG(GR3208) | 0,38 | |
| | 0CUG(GR2025) vs 123CUG(GR2024) | 0,14 | |
| | 0CUG(GR2025) vs 123CUG(GR3207) | 0,14 | |
| | 0CUG(GR3208) vs 123CUG(GR2024) | 0,49 | |
| | 0CUG(GR3208) vs 123CUG(GR3207) | 0,08 | |
| **Supplementary Figure 3B - *t*-test** | | | |
| *kat-1* | 0CUG(GR2025) vs 0CUG(GR3208) | 0,38 | n.a. |
| | 0CUG(GR2025) vs 123CUG(GR2024) | 6,43E-04 | |
| | 0CUG(GR2025) vs 123CUG(GR3207) | 0,59 | |
| | 0CUG(GR3208) vs 123CUG(GR2024) | 4,76E-04 | |
| | 0CUG(GR3208) vs 123CUG(GR3207) | 0,37 | |
| *hmgr-1* | 0CUG(GR2025) vs 0CUG(GR3208) | 0,20 | |
| | 0CUG(GR2025) vs 123CUG(GR2024) | 0,11 | |
| | 0CUG(GR2025) vs 123CUG(GR3207) | 0,21 | |
| | 0CUG(GR3208) vs 123CUG(GR2024) | 0,22 | |
| | 0CUG(GR3208) vs 123CUG(GR3207) | 0,04 | |
| *fdps-1* | 0CUG(GR2025) vs 0CUG(GR3208) | 0,96 | |
| | 0CUG(GR2025) vs 123CUG(GR2024) | 0,27 | |
| | 0CUG(GR2025) vs 123CUG(GR3207) | 0,29 | |
| | 0CUG(GR3208) vs 123CUG(GR2024) | 0,23 | |
| | 0CUG(GR3208) vs 123CUG(GR3207) | 0,27 | |
| Y48B6A.13 | 0CUG(GR2025) vs 0CUG(GR3208) | 0,43 | |
| | 0CUG(GR2025) vs 123CUG(GR2024) | 0,12 | |
| | 0CUG(GR2025) vs 123CUG(GR3207) | 0,06 | |
| | 0CUG(GR3208) vs 123CUG(GR2024) | 0,16 | |

| | | | |
|---|---|---|---|
| | 0CUG(GR3208) vs 123CUG(GR3207) | 0,03 | |
| **Supplementary Figure 3C - *t*-test** | | | |
| 0CUG(GR2025) vs 0CUG(GR3208) | | 0,56 | n.a. |
| 0CUG(GR2025) vs 123CUG(GR2024) | | 0,42 | |
| 0CUG(GR2025) vs 123CUG(GR3207) | | 0,34 | |
| 0CUG(GR3208) vs 123CUG(GR2024) | | 0,27 | |
| 0CUG(GR3208) vs 123CUG(GR3207) | | 0,26 | |
| **Supplementary Figure 3D - *t*-test** | | | |
| 0CUG(GR2025) vs 0CUG(GR3208) | | 0,29 | n.a. |
| 0CUG(GR2025) vs 123CUG(GR2024) | | 0,36 | |
| 0CUG(GR2025) vs 123CUG(GR3207) | | 0,53 | |
| 0CUG(GR3208) vs 123CUG(GR2024) | | 0,77 | |
| 0CUG(GR3208) vs 123CUG(GR3207) | | 0,87 | |
| **Supplementary Figure 5B - Wilcoxon** | | | |
| 0CUG | EV vs *gas-1* 20% | <0,0001 | p < 0,01 |
| | EV vs *nuo-6* 20% | 1,90E-03 | |
| | EV vs *mev-1* 40% | <0,0001 | |
| | EV vs *sdha-1* | <0,0001 | |
| 123CUG | EV vs *gas-1* 20% | 0,09 | |
| | EV vs *nuo-6* 20% | 0,14 | |
| | EV vs *mev-1* 40% | <0,0001 | |
| | EV vs *sdha-1* | 0,12 | |
| 0CUG EV vs 123CUG EV | | <0,0001 | |
| **Supplementary Figure 5D - 2-way Anova with bonferroni correction** | | | |
| Basal | 0CUG EV vs 0CUG *coq-1* | 0,05 | n.a. |
| | 0CUG EV vs 123CUG EV | 1,81E-17 | |
| | 123CUG EV vs 123CUG *coq-1* | 1,53E-13 | |
| FCCP | 0CUG EV vs 0CUG *coq-1* | 1,00 | |
| | 0CUG EV vs 123CUG EV | 0,01 | |
| | 123CUG EV vs 123CUG *coq-1* | 1,62E-08 | |
| **Supplementary Figure 5E - *t*-test** | | | |
| 0CUG(GR2025) vs 0CUG(GR3208) | | 0,37 | n.a. |
| 0CUG(GR2025) vs 123CUG(GR2024) | | 0,22 | |
| 0CUG(GR2025) vs 123CUG(GR3207) | | 0,07 | |
| 0CUG(GR3208) vs 123CUG(GR2024) | | 0,71 | |
| 0CUG(GR3208) vs 123CUG(GR3207) | | 0,10 | |
| **Supplementary Figure 6C - *t*-test** | | | |
| wt vs *mbl-1* | | 0,21 | n.a. |
| **Supplementary Figure 6D - *t*-test** | | | |
| wt vs *mbl-1* | | 0,76 | n.a. |
